# Neural bases of risky decisions involving nicotine vapor versus monetary reward

**DOI:** 10.1101/2021.04.14.439907

**Authors:** Priyamvada Modak, Christian Hutslar, Rebecca Polk, Emily Atkinson, Lindsey Fisher, Jon Macy, Laurie Chassin, Clark Presson, Peter R. Finn, Joshua W Brown

## Abstract

Substantial effort has gone into neuroimaging studies of neural mechanisms underlying addiction. Human studies of smoking typically either give monetary reward during an fMRI task or else allow subjects to smoke outside the scanner, after the session. This raises a fundamental issue of construct validity, as it is unclear whether the same neural mechanisms process decisions about nicotine that process decisions about money. To address this, we developed a novel MR-compatible nicotine vaping device, such that access to nicotine vapor could be controlled and monitored. We recruited heavy smokers to perform a gambling task with nicotine and monetary reward on separate days. This allowed a direct comparison between the neural mechanisms of choosing and receiving immediate drug vs. monetary reward. We found substantial differences in the neural mechanisms that underlie risky choices about money vs. drug reward, including a reversal of the well-known error effects in the medial prefrontal cortex.

## Introduction

Tobacco use remains a leading preventable cause of death in the United States and accounts for about 20% of deaths per year (CDC, health effects of cigarette smoking 2017). Nicotine addiction is sustained through both positive and negative reinforcement. Nicotine leads to positive reinforcement in form of activating rewarding pathways in the brain and, for chronic users, negative reinforcement in the form of eliminating craving and other withdrawal-related symptoms (Dani and Heinemann, 1996; McLaughlin et al., 2017; Mclaughlin et al., 2015). These effects pose a challenge for the treatment efforts to rehabilitate people dependent on nicotine. Understanding how people make decisions about using tobacco despite the well-known associated health risks is important to develop effective ways of discouraging tobacco use, and treatments for the current users wanting to quit.

The first challenge is to find appropriate paradigms to understand the decisions to use tobacco. One way to think about the decision to use tobacco is as an undervaluation of delayed outcomes where a person chooses consumption of nicotine for its immediate rewarding outcomes while ignoring more severe and detrimental long-term outcomes. Delay discounting tasks are often used to study this and find that nicotine-dependent subjects show greater discounting of monetary rewards as compared to healthy controls (Bickel et al., 1999; Dean et al., 2011; Mitchell, 1999).

Beyond delay discounting, tobacco use decisions may involve a greater tendency to take risks such that when faced with a decision to either use or not use tobacco, the person chooses the option of using, which is rewarding but also entails greater risk of negative outcomes. Various cognitive tasks have been used to study risky decision-making in general, but these suffer from potential confounds when applied to addiction. The tasks generally involve monetary reward, such as the Iowa Gambling Task (Bechara et al., 1994) and the Balloon Analogue Risk Task, or BART (Lejuez et al., 2002), in which subjects must pump up a balloon, but without popping it, to gain money. Initial studies of smokers with the BART had shown the smokers more willing to take risks (Lejuez et al., 2005, 2003), but this is disputed (Dean et al., 2011). In the BART, the option of cashing-in leads to immediate reward and end of the current trial while the option of inflation delays the reward and lengthens the current trial. It is also possible that smokers cashed in earlier (leading to lower mean adjusted pumps) because they discounted the delayed reward. Thus, delay discounting may be confounded with risk avoidance.

In addition to task confounds, there is the issue of construct validity in studying decisions about drug usage – it is unclear that studies involving monetary reward tap the same neural mechanisms involved in drug reward, or even that monetary reward processing remains intact. Studies using task-based fMRI in nicotine dependence generally use non-drug rewards, like money, to study decision making in nicotine-dependent subjects (Alexander et al., 2015; Dean et al., 2011; Fukunaga et al., 2013; Galván et al., 2013). However, research has shown that there is altered sensitivity to monetary rewards in substance use. In cocaine users, almost half of the subjects with cocaine addiction showed significantly reduced subjective sensitivity to monetary rewards as compared to healthy controls (Goldstein et al., 2007). It is possible that the subjects had a reduced sensitivity to monetary rewards prior to the development of cocaine addiction and not because of it. In this respect, Just et al. (2019) showed altered connectivity of putamen with frontal regions in participants with familial risk of drug dependence irrespective of drug usage during monetary incentive delay task. Furthermore, the activation of mesocorticolimbic structures to anticipating monetary rewards relative to anticipating cigarette rewards is reduced in dependent smokers as compared to occasional smokers, and motivation to earn monetary rewards relative to cigarette rewards is also reduced in dependent smokers vs. occasional smokers (Bühler et al., 2010). Dependent smokers are more willing to pay for cigarettes as compared to monetary vouchers whereas occasional smokers are more willing to pay for vouchers than cigarettes when the vouchers and cigarettes were matched in their monetary worth (Lawn et al., 2020). Nicotine abstinence in dependent smokers can also reduce monetary reward anticipation-related activity in the bilateral caudate and middle prefrontal cortex (Sweitzer et al., 2014). Further, the valence-related activity for monetary rewards in the anterior cingulate cortex (ACC) in Rose et al. (2013) was negatively correlated with smoking severity (cigarettes per day). These studies collectively show that the sensitivity to monetary or non-drug rewards and cigarette rewards in smokers is different, suggesting that monetary or non-drug rewards cannot be used as a proxy to study decisions about nicotine consumption in nicotine-dependent subjects. In the present work, we have compared the choice behavior and neural underpinnings of decisions about both nicotine and money to better understand the utility and limitations of non-drug rewards in nicotine dependence.

There are previous task-based functional Magnetic Resonance Imaging (fMRI) studies that have used nicotine as a reward, however, the reward was in form of letting the subjects smoke after the task, outside the scanner (Lawn et al., 2020; MacKillop et al., 2012). However, smokers display greater discounting of money than controls, and they discount cigarettes even more than the money (Bickel et al., 1999), but this has not always been replicated (MacKillop et al., 2012). Likewise, e-cigarette users show greater discounting of e-cigarettes as compared to money (Pericot-Valverde et al., 2020). Moreover, Wilson et al. (2005) have shown that activation to smoking cues in smokers is different depending on the smoking expectancy, that is whether they could smoke immediately after the task or after a delay of an hour. Providing earned nicotine rewards to the subjects only after the task session might lead to confounding by the effects of delay discounting of the rewards, and also a confounding effect of smoking expectancy on the choice behavior and cognition of the smokers. To avoid these potential confounds, we developed a novel method to provide nicotine rewards inside the scanner, immediately after the subjects won at the task, in form of nicotine vapor from a custom e-cigarette.

Here we employ a novel gambling task that orthogonalizes the gamble characteristics: expected value, probability of loss, and variance of the gamble outcomes. This allows us to separately evaluate the effects of reward value as well as the two conceptualizations of risk: the probability of loss and variance of gamble outcomes, on both choice behavior and blood oxygenation level dependent (BOLD) activation in the brain. The task involved choosing between a gamble with three possible outcomes and a sure thing with only one outcome which was equal to certainty equivalent of the gamble. The subjects received the reward based on the outcome of this choice. We had the nicotine-dependent subjects perform this task inside the scanner on two separate days, once with nicotine as the reward and once with money as the reward. In the case of nicotine, a modified e-cigarette was used to deliver nicotine vapor inside the scanner. We addressed the following questions: 1. How do nicotine-dependent individuals make decisions about nicotine, in terms of choice behavior and neural correlates of choice parameters (expected value, probability of loss, and variance) of the gamble outcomes? 2. Can the results from decisions about money rewards in nicotine-dependent individuals be extrapolated to infer decisions about nicotine? We compared the results from money and nicotine sessions to answer this question. By delivering drug reward immediately in the scanner, we both increase the construct validity and avoid potential confounds with delay discounting.

## Results

### Behavioral Analyses

The subjective value of a gamble was mainly accounted for by its expected value, for both nicotine and money rewards, suggesting that the subjects were motivated to gain reward. The average certainty equivalent (CE) values of all subjects for each of the five gambles (such that we had five values from each subject corresponding to the CE of five gambles) were regressed with the gamble characteristics, namely, probability of loss, variance, and expected value. As shown in table 1, for both money and nicotine sessions, the effect of the expected value of gamble on the CE was positive and significant (Nicotine: b = 0.92, p = 0.006; Money: b = 1.001, p<0.001) indicating greater CE and thus, greater likelihood of choosing the gamble when the expected value of a gamble was greater. These effects remained significant even after Bonferroni correction was performed. The effects of other gamble characteristics were not significant. The result of the effect of expected value supports the assumption that subjects wanted the rewards and were not averse to nicotine or money during the sessions. This result helps us rule out the possibility that subjects were satiated and did not want nicotine or were not engaged in the task. We did however find a subset of subjects who appeared to be unmotivated in the task, but the rest of our results were not significantly different with those subjects excluded, so we include them in the following analyses (Supplementary material, section E). We also found associations among initial CO, as a measure of nicotine satiety, and gamble preferences (Supplementary material, section F).

**Table 1:** Regression coefficients of probability of loss, variance and expected value of gambles on the certainty equivalent values of the gambles, indicating the association of each factor with relative preference for the gamble.

|  | Money |  | Nicotine |  |
| --- | --- | --- | --- | --- |
|  | Beta weight | p value | Beta weight | p value |
| **Constant** | 0.074 | 0.778 | -0.14 | 0.85 |
| **Ploss** | -0.351 | 0.41 | -0.43 | 0.71 |
| **Var** | 0.316 | 0.073 | 0.92 | 0.055 |
| **EV** | 0.975* | <0.001 | 0.92* | 0.006 |

Ploss – Probability of loss, Var – Variance, EV – Expected value

**Figure 1:**
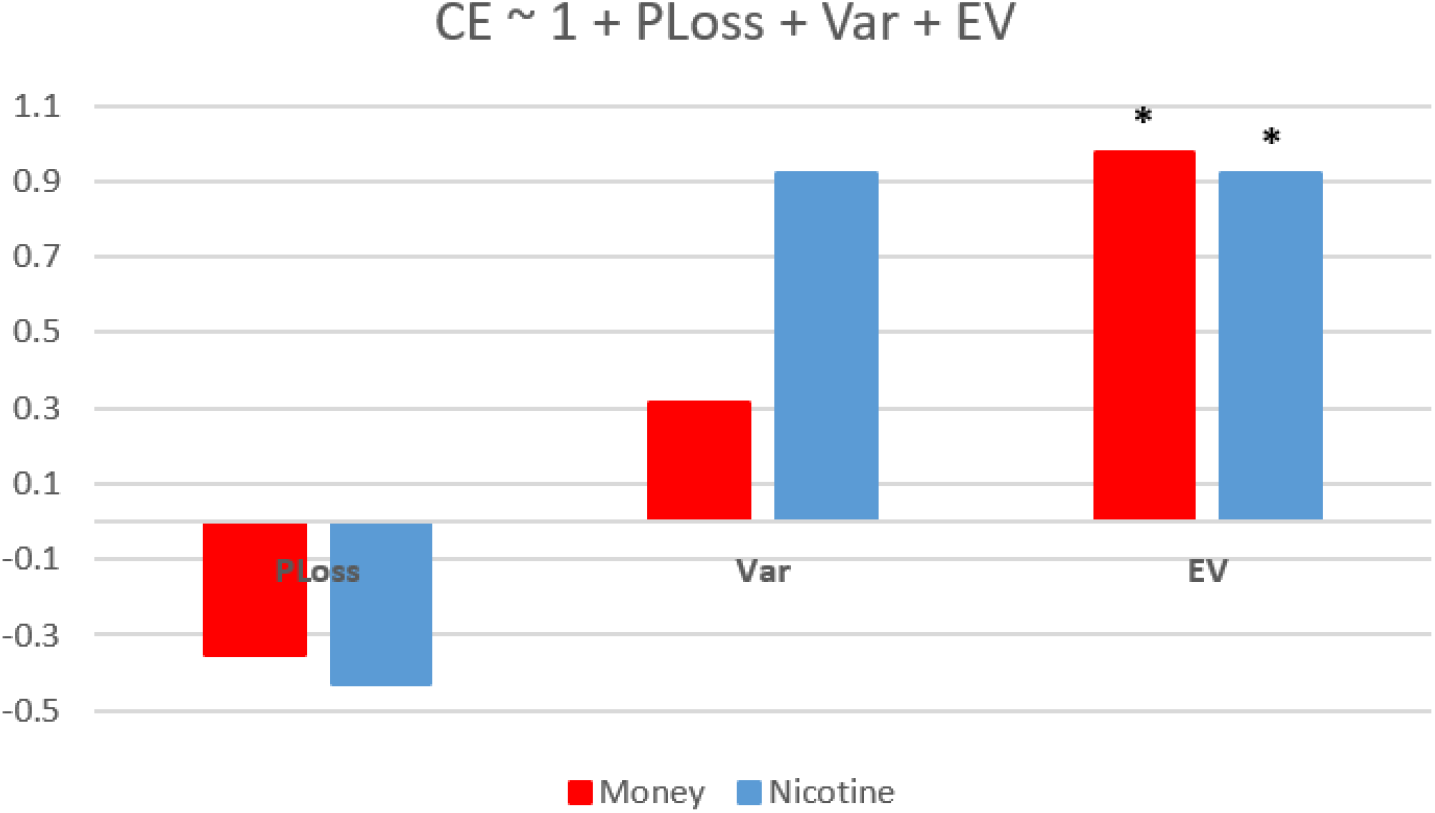
Effects of gamble characteristics on certainty equivalent of the gambles in money and nicotine sessions.

### fMRI analyses

#### Effect of gamble characteristics during choice phase – Nicotine

We investigated the BOLD effects of risk and value of nicotine, and we found evidence of nicotine reward anticipation during the choice phase of each trial. As shown in figure 2, we found that the anterior cingulate cortex and caudate show a negative loading on the probability of loss of nicotine, which indicates a positive loading on the probability of winning nicotine (peak voxel MNI: 18, -14, 28). The observed effect in these regions might reflect reward anticipation in case of nicotine reward. We also observed a region in the left occipital lobe (peak voxel MNI: -18, -94, 16) showing a negative loading on the nicotine gamble variance regressor. Refer to supplementary tables S7, and S10 for additional information about these clusters.

**Figure 2:**
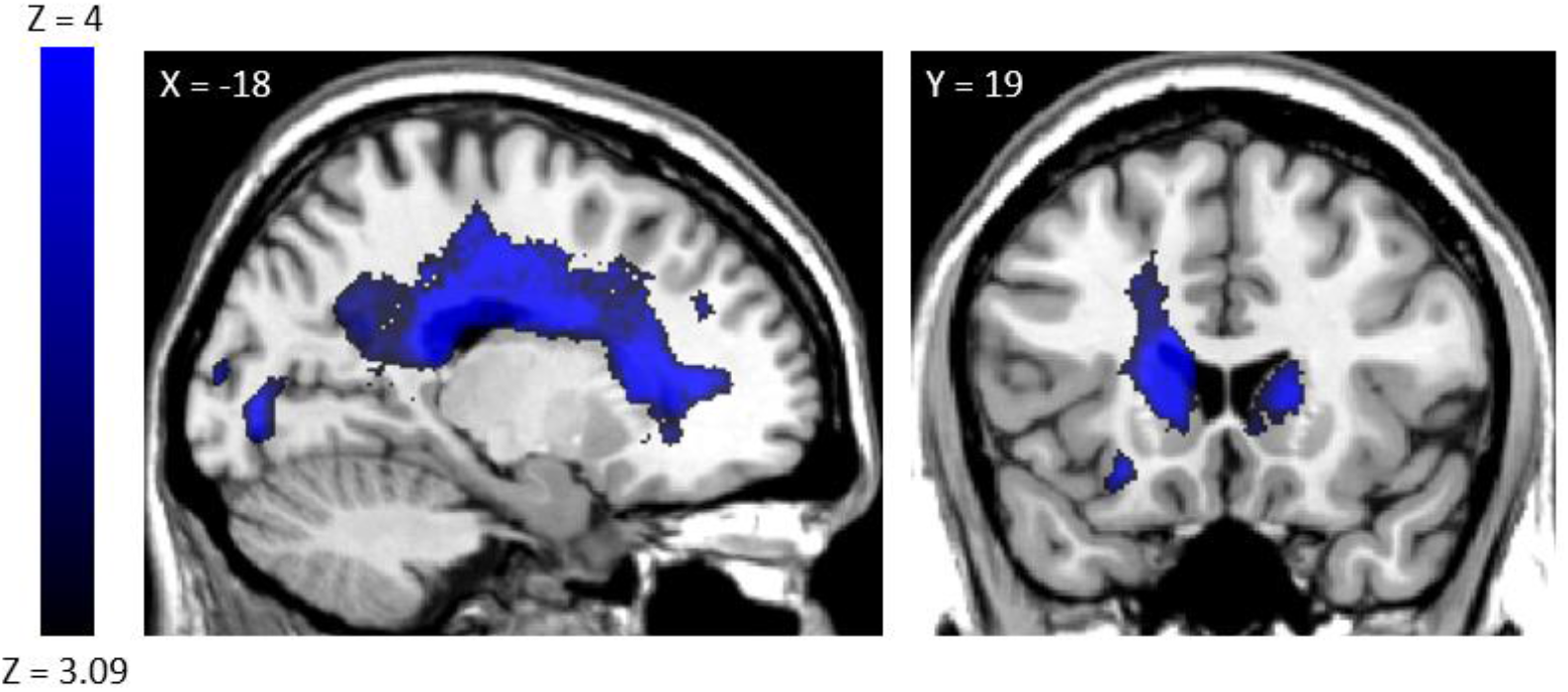
Nicotine decision effects. The cingulate cortex and caudate show activity associated with probability of winning nicotine when the gamble was chosen. Regions shown with negative loading on probability of loss (blue) regressors in nicotine session. Left: Bilateral cingulate. Right: Bilateral caudate

We did not observe any regions that showed a significant positive effect of the probability of loss, variance, and expected value regressors in nicotine sessions. It is possible that with a bigger sample size we might be able to identify regions that show these effects. The lack of effects here is not due to issues with our methods, because we were able to find such effects in the case of monetary rewards, as we show below.

#### Effect of gamble characteristics during choice phase – Money

We did not observe the same regions to show reward anticipation (i.e. effect of the probability of winning) in the money session, despite strong effects in the nicotine session. The regions that showed the effects of the probability of loss, variance, and expected value in the money session are listed in tables S5, S8, and S11 in the supplementary material. There were regions in middle/superior frontal gyrus (peak voxel MNI: -10, 38, 40; 18, 40, 40; -16, 46, 22) and middle temporal gyrus (MTG) (peak voxel MNI: -46, -36, -12; -46, -56, 20; 46, -44, 2; -52, -54, -4) that showed overlapping effects of the probability of loss and variance in the money session, as shown in figure 3.

**Figure 3:**
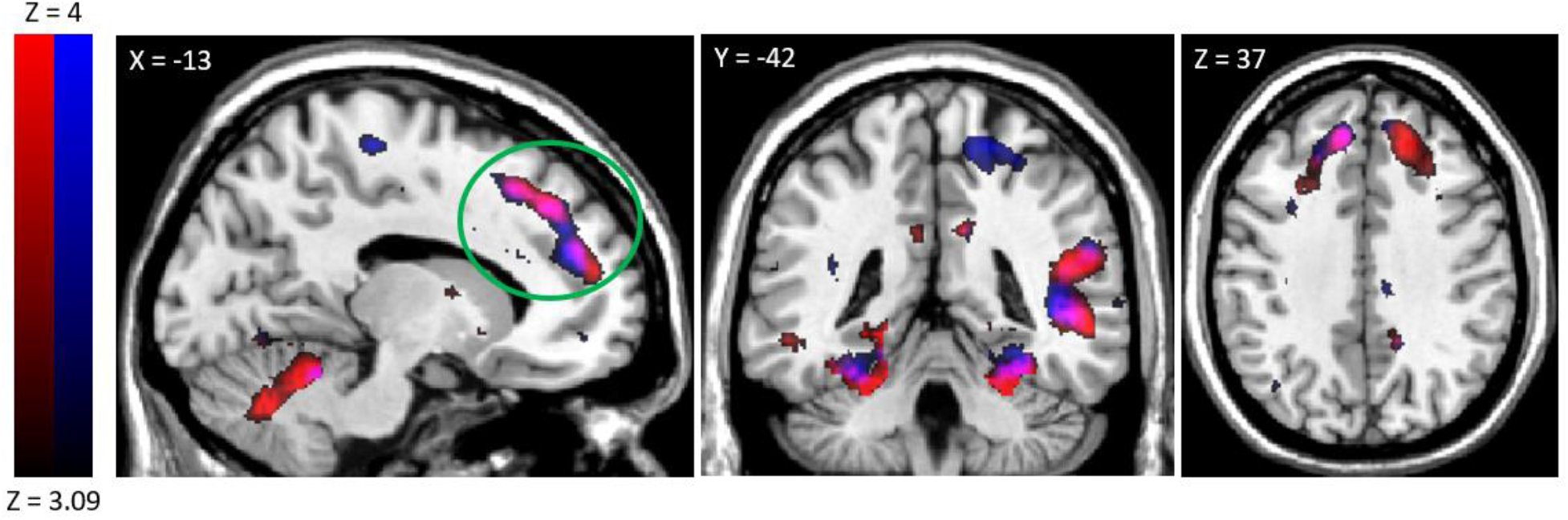
Monetary decision effects. Regions with positive loading on both variance (red) and probability of loss (blue) regressors in money session. Left & middle: Middle/Superior Frontal Gyrus ((Circled in solid green line in middle). Right: Middle Temporal Gyrus (Circled in solid green line). Circled clusters are significant at cluster p < 0.05.

**Figure 4:**
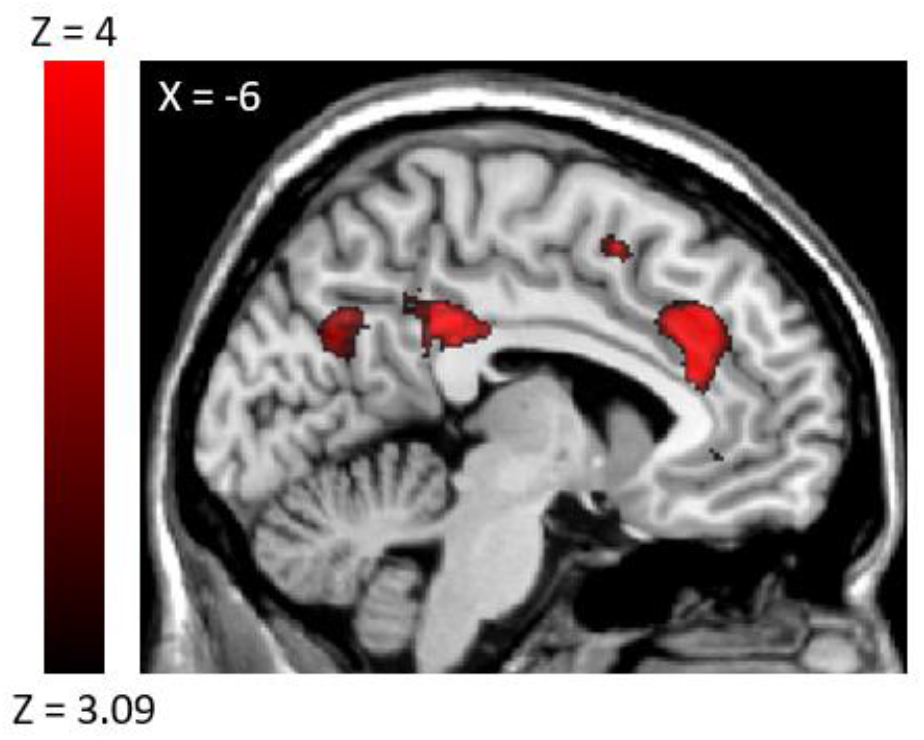
Positive loading on the sure thing value in money sessions

#### Effect of sure thing value during choice phase

The effect of sure thing value during choice phase in money session was observed in anterior (peak voxel MNI: -6, 32, 32) and posterior (peak voxel MNI: -4, -36, 34) cingulate cortex in the money sessions. No significant effect was observed in the case of nicotine sessions. Refer to table S22 for additional information about these clusters.

#### Effects of winning and losing during feedback phase in money and nicotine sessions

The effects observed in the feedback phase of the session where the subjects won (i.e., received a non-zero reward) or lost (i.e., received zero rewards) were particularly noteworthy (figure 5), as they show a striking reversal of the error effects. There were overlapping yet separate regions spanning the cingulate gyrus and parts of the middle and superior frontal gyrus (MFG, SFG) that showed significantly higher activation to money loss as compared to money win (peak voxel MNI: 14, 8, 62; 42, 16, 0; -32, 16, -10) while the opposite was true for nicotine – there was significantly higher activation to nicotine win as compared to nicotine loss (peak voxel MNI: -36, -6, 12). We refer to this as the error reversal effect from now onwards. There was also a small region in BA 32 (ACC) (peak voxel MNI: 4, 28, 28) that shows the effect of self-reported craving on its relative activation to nicotine win, such that higher self-reported craving was associated with lower relative activation of this region for nicotine win as compared to nicotine loss. For generating the plots in figure 5B, we found the activation of parts of red and blue clusters in the cingulate gyrus.

**Figure 5:**
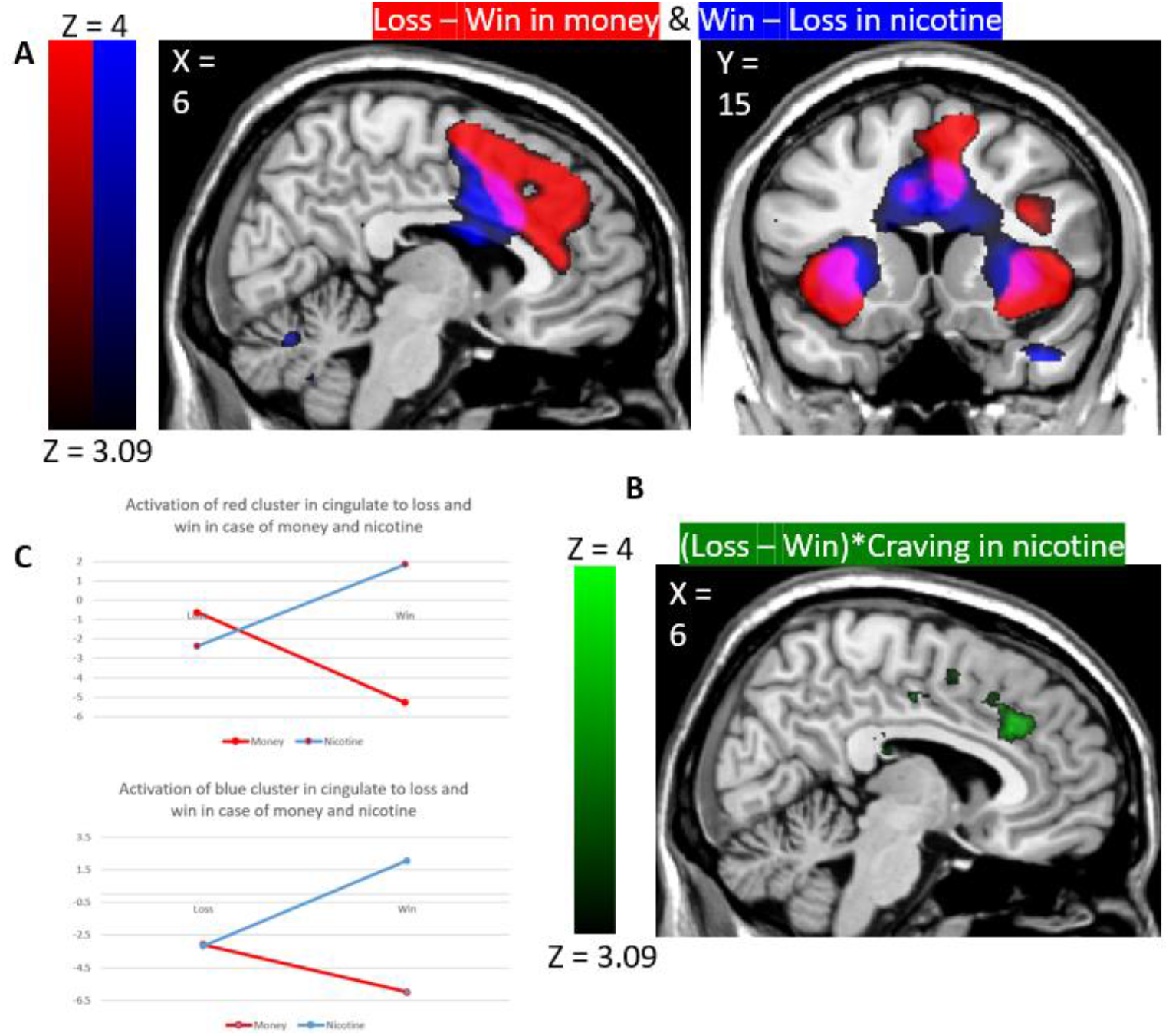
Feedback effect reversal in money vs nicotine session. A: Regions that load significantly more on money loss than money win are shown in red. Regions that load significantly more on nicotine win than nicotine loss are shown in blue. B: Regions that show interaction of loss – win in nicotine session and self-reported craving are shown in green. C: Graphs showing activation of red and blue clusters in cingulate to loss and win in case of both, money and nicotine.

#### Effect of inhaling nicotine vapor

We found a range of motor cortical activity and functionally connected regions involved with inhaling nicotine. In the money sessions, each trial ended after the feedback phase, and the next trial began after a jitter. However, in the nicotine session, the feedback phase was followed by inhalation for the duration equal to the gamble outcome in seconds (figure 8). As expected, the motor cortex area for lips/mouth showed the effect of inhalation event, presumably reflecting mouth movements (peak voxel MNI: 68, -8, 12; -64, -12, 14). This can be seen in figure 6 (with more details in Tables S20 and S21 in the supplementary material).

**Figure 6:**
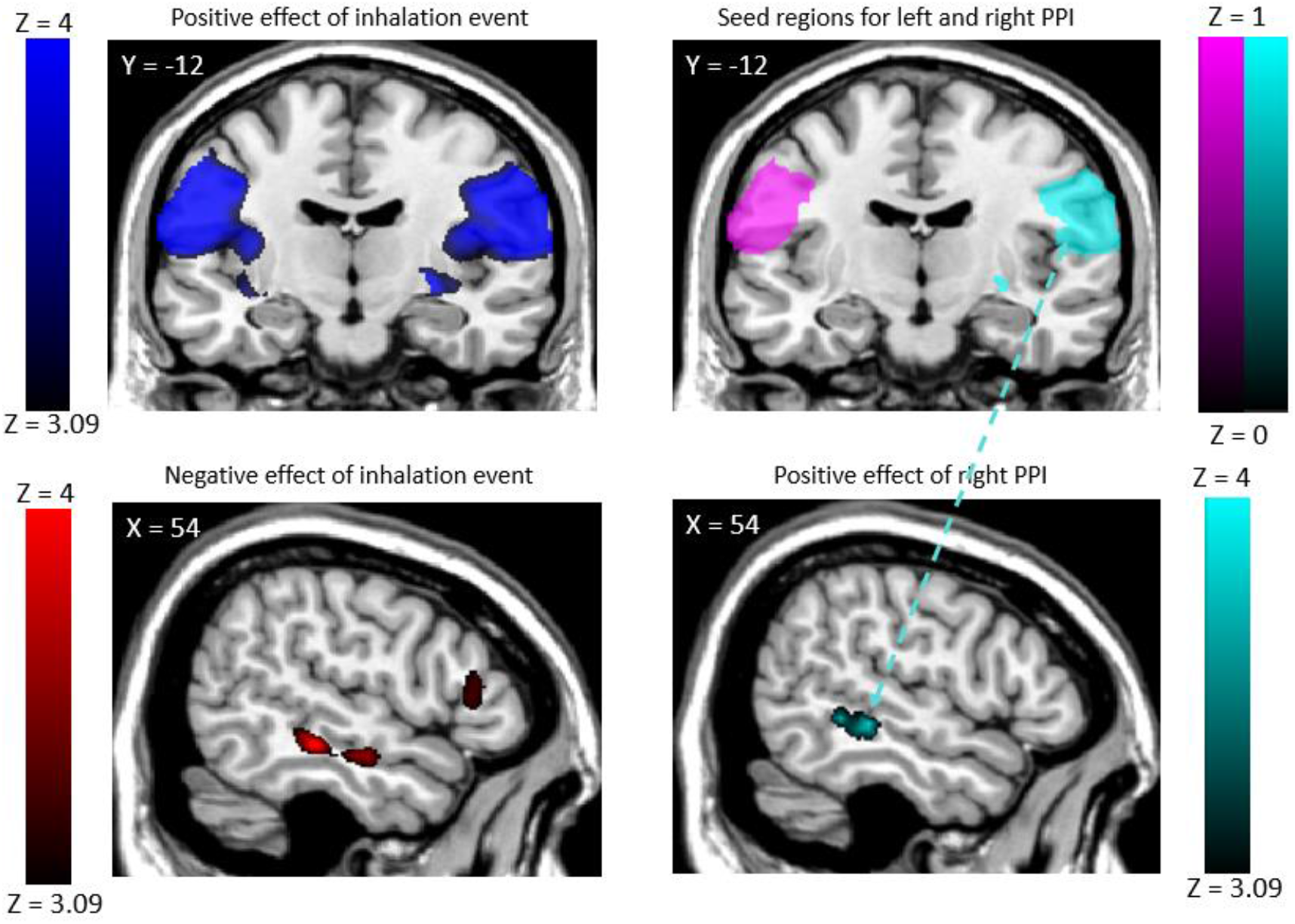
Psychophysiological interaction with motor regions. Top-left:Positive loading of bilateral motor cortex on inhalation event regressor.Top-right: Seed regions for left and right PPI analysis for individual subjects were identified within these regions using small volume correction. Bottom-left: Negative loading of bilateral MTL on inhalation event regressor. Bottom-right: Positive loading of right Middle Temporal Lobe (MTL) on right PPI regressor.

**Figure 7:**
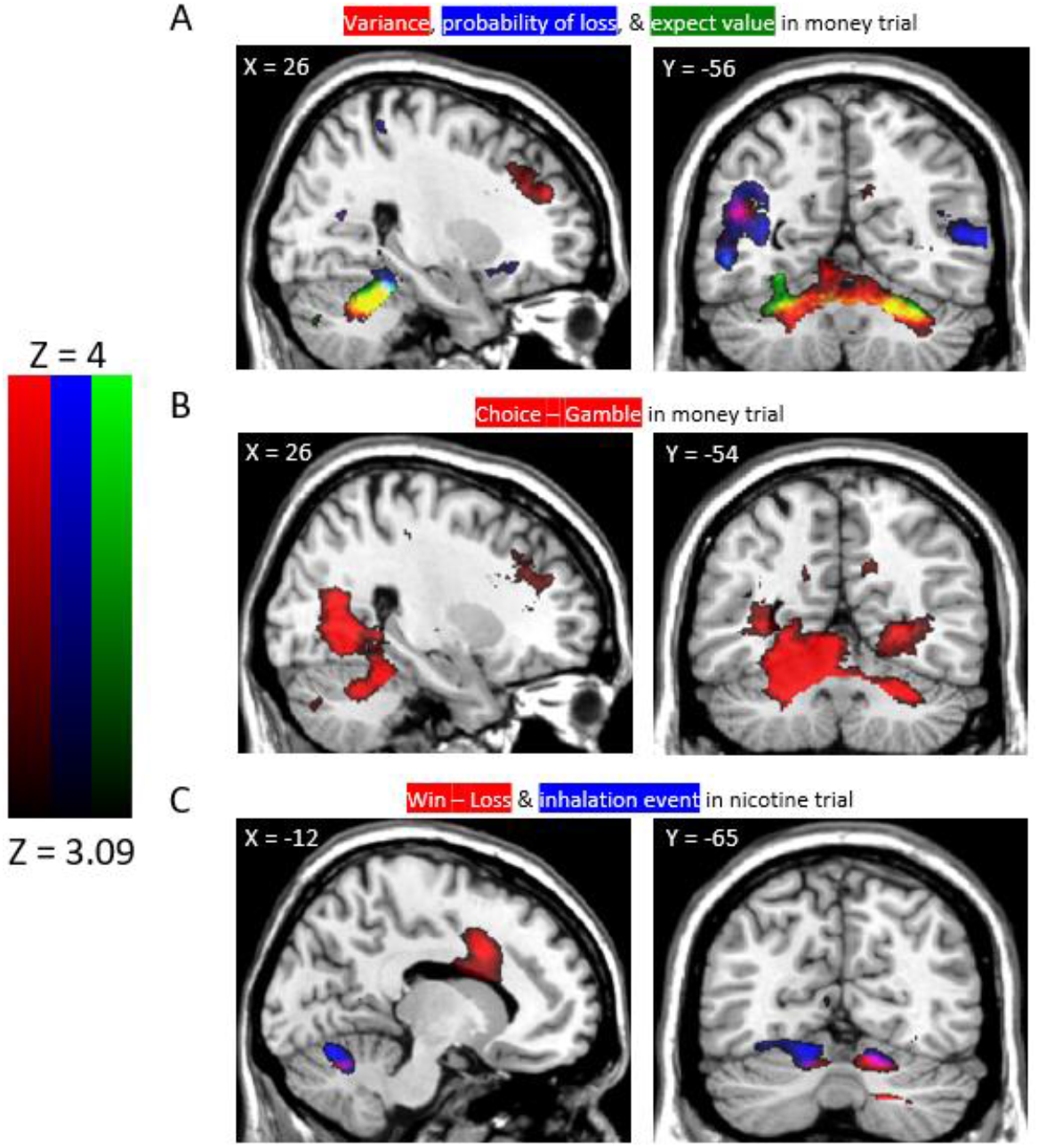
Task effects in cerebellum. Rows A and B show effects in cerebellum during choice. Row C shows effects during feedback. A: Effects of variance (red), probability of loss (blue), and expected value (green) of gamble in money session; B: Effect of choice – gamble in money session (red); C: Effect of win – loss (red) and inhalation event (blue) in nicotine session

**Figure 8:**
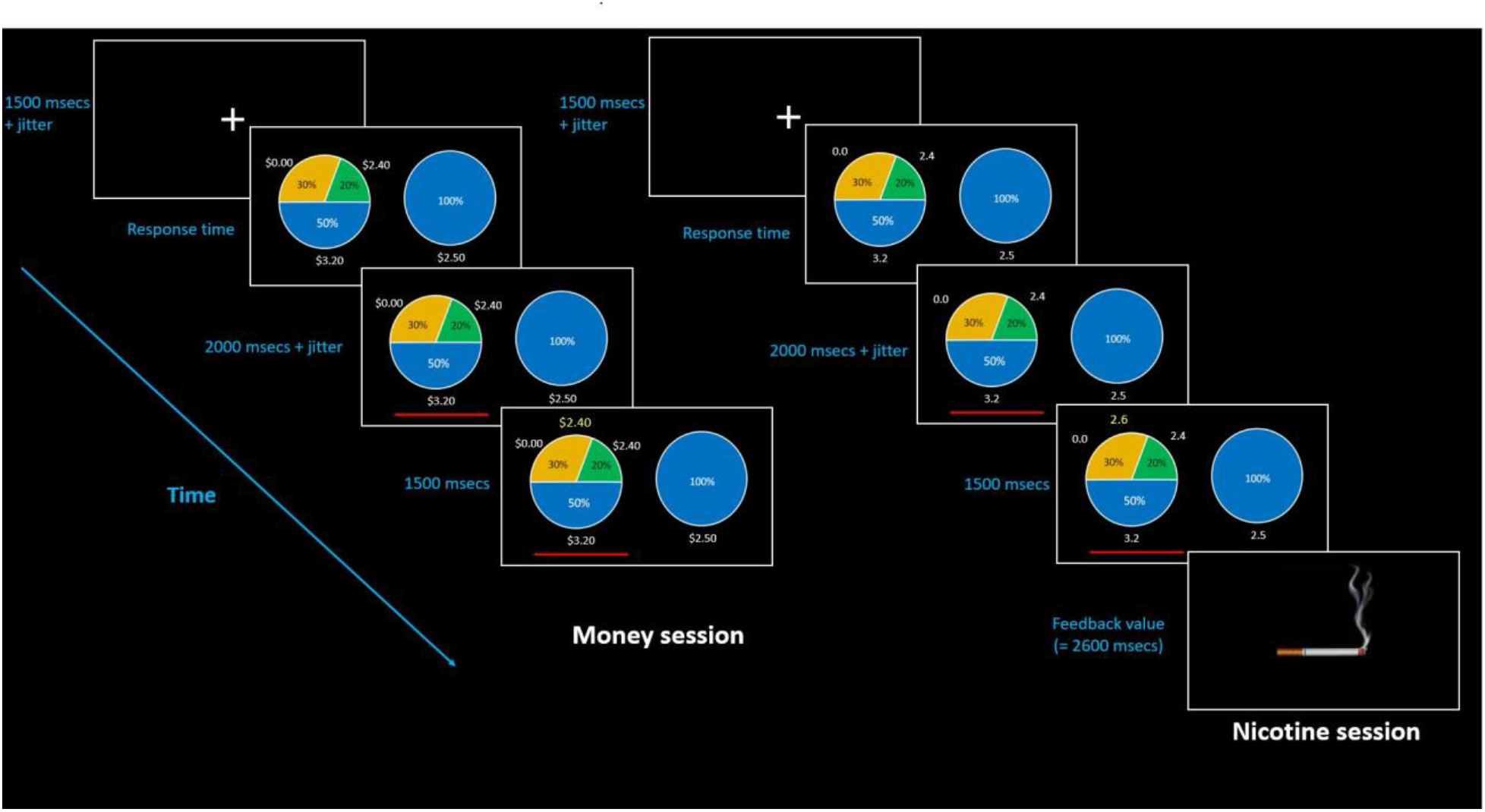
Task time courses. Each trial begins with a fixation period, after which the choice is presented. Immediately following a response, the chosen option is underlined in red, but the outcome is not revealed until after a delay. This affords separate estimation of the BOLD responses to choice vs. feedback about the outcome. In the nicotine trials, there is additionally a time for nicotine vapor inhalation after the feedback is first revealed.

We were further interested to find which regions were indirectly involved in the inhalation of nicotine. In this respect, we performed psychophysiological interaction (PPI) analysis with the motor regions that showed significant activation in inhalation events as the seed regions. Right MTG (peak voxel MNI 52, -42, -2) showed an association with the right seed region regressor. This implies that the functional connectivity between the right MTG and right motor cortex increases during the inhalation event.

Moreover, the main effect of inhalation on bilateral MTG is negative, that is, it shows deactivation upon inhalation in nicotine session (peak voxel MNI: -58, -28, -6 ;54, -30, -10). This means that the absolute activation of right MTG went down when there was inhalation but the trend in the changes in its activity matched the trends in the activity of the right motor cortex, more closely during inhalation than at other times.

Please refer to tables S33, S34, and S35 for all the results of PPI analyses.

#### Cerebellar effects

Many task effects were observed in the cerebellum. As shown in figure 7, regions in the cerebellum showed a positive effect of the probability of loss (peak voxel MNI: -30, -40, -22; 28, -40, -20), variance (peak voxel MNI: -28, -50, -28), the expected value of gamble (peak voxel MNI: -26, -48, -26), and the event of choosing the sure thing (peak voxel MNI: -20, -48, -24), in money sessions. The cerebellum also showed a positive effect of win – loss (peak voxel MNI: 14, -66, -22), and the event of inhalation (peak voxel MNI: -12, -68, -16) in nicotine sessions. Interestingly, the effects in the cerebellum in the money sessions were in the culmen, but in nicotine sessions they were found in the declive. Refer to table S1, S5, S8, S11, S17, and S20 for additional information about these clusters, and to section G in supplementary material for a discussion of the cerebellar effects.

Please refer to section H in supplementary material for results from the analysis of money session data from only 20 subjects that participated in both, money, and nicotine sessions. Although the task effects are weaker in this analysis as expected, we still observe error reversal between money and nicotine sessions. This rules-out the possibility that merely the difference in number of participants between money and nicotine sessions could have caused the observed error reversal.

## Discussion

As a whole, our results show that to understand the neural basis of decision-making in addiction, it is not sufficient to study only decision-making tasks with monetary reward. There are a number of differences in the neural responses to choices involving money vs. drug reward, and these have not been explored extensively in the literature previously. There is only one study, to the best of our knowledge, which had subjects consuming nicotine inside the fMRI scanner (Wall et al., 2017). However, that study consisted of ad-libitum and cued smoking inside the scanner and was not designed to inform about the choice behavior in terms of risk-taking tendencies and the neural correlates of different aspects of decision making about nicotine in nicotine dependent subjects. There is one more study where the substance of choice was consumed inside the scanner, however, that was with cocaine-dependent subjects (Risinger et al., 2005).

### Behavioral results

The expected value of the gambles was the best predictor of the choice behavior such that the greater expected value of the gamble led to a greater tendency to choose the gamble. This confirms that the subjects wanted the rewards and were engaged in the task. The negative correlation between beta weights on expected value and initial CO is consistent with the idea that subjects who were sated with nicotine were in turn less likely to pursue the highest expected value of nicotine. The positive correlation between beta weights on variance and initial CO reflects that subjects with higher initial CO were more willing to take risks by choosing the gamble with the greater variance.

### Effects during the choice phase in nicotine and money sessions

We found several regions including MFG, SFG, MTG, and regions around the cingulate cortex showing positive loading on the probability of loss and variance in the case of monetary reward. This is consistent with previous studies with monetary rewards, implicating these regions in decision making, specifically in the representation of risk (Fukunaga et al., 2018), in option evaluation, and outcome evaluation in nicotine dependence (Lawn et al., 2020; MacKillop et al., 2012; Rose et al., 2013), in other substance dependent populations (Cservenka and Nagel, 2012; Gelskov et al., 2016; Paulus et al., 2005) as well as in healthy samples (Alexander and Brown, 2014; Ernst et al., 2002; Fukunaga et al., 2018; Jahn et al., 2014; Krain et al., 2006).

The task that was used in Fukunaga et al. (2018) most closely matches the task we have used here. The results from that study revealed a strong effect of variance in right MFG/IFG and cingulate cortex. We have also observed the effect of money variance in these regions. The size of the regions showing the effect is however smaller in the current study. The present study differs from the previous one (Fukunaga et al., 2018) in that the five gambles used in the current study were constructed to orthogonalize probability of loss, variance, and the expected value, but the previous study orthogonalized probability of loss, variance and maximum possible loss. In particular, the previous study did not manipulate the expected values across gambles as we have done here. The difference in the extent of the loading on gamble variance in both studies can be attributed to the difference in subject population between both the studies. In our study, the subject population consists of nicotine-dependent individuals who were asked to remain abstinent for at least 6 hours prior to the experiment while the previous study used a random sample of individuals which may or may not have been abusing drugs. Therefore, the observed differences between activation of regions could stem from the drug-induced neuroadaptations in the subject population in this study. This is consistent with previous work showing a negative correlation of resting-state functional connectivity between dorsal ACC and striatum with the severity of nicotine dependence (Hong et al., 2009), and a reduction in monetary reward sensitivity in cocaine dependence (Goldstein et al., 2007). The ACC in particular has shown reduced sensitivity to monetary valence in abstinent smokers (Rose et al., 2013), consistent with our findings here.

We found BOLD correlates of the expected value of the gamble and sure thing options in the money sessions. A positive loading of the BOLD signal on the expected value of gamble was found in the middle temporal lobe and cerebellum (Table S11), but surprisingly, we found no effect of expected value at the time of choice in the vmPFC, OFC, or nucleus accumbens (and not in nicotine trials either). This is noteworthy given earlier findings of value-related activity in these areas, both for money and primary (juice) reward (Kim et al., 2011). This is unlikely to reflect signal dropout in the OFC area, as we have found effects in these regions in previous studies with nearly identical scan protocols (Fukunaga et al., 2012). A positive loading on sure-thing value was observed in the anterior and posterior cingulate cortex (Figure 4). This sure-thing loading implies that these regions could be representing the value of a less risky option and furthermore might be involved in driving the choice behavior against taking the risk. To investigate this, we looked at the correlation between the effect of sure thing value in these regions and the certainty equivalent values across all the subjects in the money session. The correlation was not significant but was negative, so we cannot make a definite conclusion. This does not rule out a role for the ACC in risk avoidance (Brown and Braver, 2007), or seeking a sure reward. Further, ACC has been implicated in representing regret (Coricelli et al., 2005), thus, the effect of sure-thing value in anterior cingulate could also be signaling anticipated regret related to choosing the gamble. To evaluate this possibility, we compared the effect of sure thing value in events when the gamble was chosen vs. events when the sure thing was chosen. We reasoned that if the cingulate was indeed signaling anticipated regret, the activation should be more when gamble was chosen. We did not find any significant difference between the effect of sure-thing in the two events.

In the nicotine sessions, we observed the effect of the probability of winning in the right cingulate gyrus and left caudate. Previous studies have shown the effect of monetary reward expectancy on these regions in smokers (Rose et al., 2013) and in pathological gamblers (Gelskov et al., 2016). Our study now demonstrates this effect directly in anticipation of nicotine rewards in smokers. We did not find the effect of the expected value of the gamble in any of the regions in nicotine sessions, although we did see a cluster in the ventral medial prefrontal cortex (vmPFC), which did not pass the cluster correction. Consistent with this, the effect of the subjective value of cigarettes in a purchase task was observed in vmPFC in Lawn et al. (2020).

### Effects in feedback phase and different effects in money vs nicotine sessions

We found a number of effects of feedback regarding the gamble outcome, for both money and nicotine rewards. The feedback effects observed in parts of the cingulate cortex, MFG, MTG, and Insula are consistent with previous studies with monetary rewards showing the effects of outcome evaluation in the same regions (Akaishi et al., 2016; Fukunaga et al., 2018; Jahn et al., 2014). Remarkably, with monetary reward, these regions showed the effect of greater activity for loss than win while in the case of nicotine reward, the regions showed the opposite effect of greater activity for win than loss. The finding represents a striking reversal of the error effects typically seen with cognitive tasks and monetary reward (Gemba et al., 1986) and challenges the long-held view of the medial PFC as an error detector (Gehring et al., 1993). It is also difficult to account for this effect in terms of surprise or prediction error (Alexander and Brown, 2011), as the task structure and outcome probabilities are the same across money and nicotine here. One possible interpretation of the greater activation for winning in nicotine trials is in terms of motivation as represented by the ACC (Holroyd and Yeung, 2012; Parvizi et al., 2013; Walton et al., 2002) – subjects may be more willing to exert effort to obtain nicotine when the nicotine is actually available.

From another perspective, it is not surprising that the feedback effects are different in money and nicotine sessions, because the effects earlier at the time of choice are also different. Also, previous research has shown that nicotine-dependent subjects have altered relative sensitivity to money and cigarettes such that the relative sensitivity to cigarettes is increased as compared to healthy controls. This has been demonstrated both behaviorally (Bühler et al., 2010; Lawn et al., 2020), and neurally in terms of the regions involved in processing both the rewards (Buhler et al., 2010; Lawn et al. 2018; Rose et al., 2013; Sweitzer et al., 2014).

There are other differences between money and nicotine sessions that might play a role in the different feedback effects, as we describe below.

### Craving

First, there were differences in craving. The self-reported craving of subjects in the money session was significantly more than the self-reported craving of subjects in nicotine session, which is not surprising as subjects were only given nicotine during the nicotine sessions and thus would not be sated during the money sessions.

The greater activity due to win feedback vs. loss feedback in nicotine trials could also be due of the win feedback acting as a drug cue, thus resulting in craving-related activation. Several studies have shown greater activation of ACC and mPFC when viewing smoking cues as compared to neutral cues (Hartwell et al., 2011; Wilson et al., 2005). Wall et al. (2017) showed increased activation of Insula and dorsal ACC (dACC) in cued smoking as compared to ad-libitum smoking. In this respect, we also found a region in the cingulate cortex that showed an interaction of the effect of self-reported craving and loss minus win in the nicotine sessions; however, this was only a small region. In the end, drug cue-related activation, craving, motivation for drug consumption may all reflect a single cued motivational construct.

Nevertheless, the valence of this motivational construct may have been different between money and nicotine sessions. The subjects might have different motivational perceptions in the money and nicotine session such that craving was acting as a ‘resource’ in nicotine session while it was acting as a ‘threat’ in money session (due to the unavailability of nicotine), which might have led to observed differences between both the sessions (Frings et al., 2015).

The greater activation of the cingulate and the parts of the frontal and temporal cortex showing error reversal effect in the case of monetary loss as compared to monetary win could be due to stress-induced craving due to the loss event specifically (Erblich et al., 2004; Hartwell et al., 2013; van Hedger et al., 2020). By the same reasoning though, the loss event in nicotine session could also lead to stress-induced craving, so it is less obvious why the effects should be different between money and nicotine reward. One possible resolution of this conundrum is that previous research shows that cue-induced craving, which could be caused by the win event in the nicotine session is more than the stress induced craving (Colamussi et al., 2007). This prior work suggests how the increased craving in these regions can explain the greater activation in monetary loss than win. Still, when we explicitly modeled self-reported craving as a regressor in the GLM, we found no main effects at the time of choice. Craving only loaded on activity at the time of feedback.

### Timing of reward

Second, the immediacy of the reward was different. The reward in the money session was not immediately consumable while the reward in the nicotine session was nicotine vapor and was immediately consumable.

The timing of nicotine reward, including its immediate availability, may have influenced the neural response. Specifically, the activation of regions, MFG, and SFG in nicotine sessions might have been reduced due to immediate reward availability (Wilson et al., 2005). In a future study, the potential effects of reward expectancy could be removed by using a consumable non-drug reward such as juice or chocolate, instead of money. However, in this study, we wanted to evaluate the utility of monetary rewards which are extensively used in substance dependence research, as a proxy for decisions about nicotine, and thus, it was necessary to use monetary rewards.

### Prediction error

We also considered whether the feedback effects may reflect a kind of positive prediction error. Specifically, it is also possible that these regions were involved in representing relief of winning the nicotine, in which the actual win is greater than the counterfactual sure thing they would have won, had they chosen the sure thing (Coricelli et al., 2005). We formed another glm (See glm D in supplementary material) to evaluate the effect of regret/relief on various regions in the feedback phase. We observed the effect of relief (opposite of regret which happens when the amount won by choosing gamble is more than sure thing value) in MTG and Insula suggesting that the greater activation to winning nicotine can be a result of relief.

### Pharmacological effects

Moreover, the error reversal observed between nicotine and money session could also stem from the fact that there was greater plasma nicotine in nicotine session, which would be expected to both reduce craving and increase the sensitivity to monetary rewards (Rose et al., 2013). In particular, previous work found that the activation of left cingulate and MFG corresponding to the successful minus unsuccessful reward feedback was observed in smokers when they were given a nicotine patch but not when they were given a placebo patch (Rose et al., 2013). Thus, it is possible that nicotine shifts the neural response to be greater in response to reward than punishment, as a pharmacological rather than cognitive effect on neural activity.

Also, greater plasma nicotine is shown to be associated with reduced activity in the NAc and putamen (Rose et al., 2013). This could lead to different reward responses in nicotine vs. money sessions due to different levels of plasma nicotine, but we find this unlikely because we controlled for plasma nicotine effects in our GLM using plasma nicotine as a nuisance regressor, as estimated from airflow using our PBPK model (Vélez de Mendizábal et al., 2015).

### Effect at the time of nicotine inhalation

Activation of motor regions during inhalation was expected since inhalation involved the movement of the mouth, at least the lips, and the lungs. From the PPI analysis, we found that the right and left MTG deactivated during inhalation, but the functional connectivity of the right MTG with the part of the right motor cortex involved in inhalation increases during inhalation. MTG has shown other task effects, as shown in figure S1 and described in section A of supplementary material, which suggests that it might be involved in the predictive and evaluative phase of the task. The implication of MTG in decision making and representing risk has been shown previously (Krain et al., 2006). Our results also show the effect of the variance of gamble outcomes and the probability of loss in the right MTG. With respect to its involvement in studies about substance dependence Gelskov et al. (2016) show an increase in activity in right MTG with an increase in gain/loss ratio of gambles in a task to reject or accept the presented gamble. We also observe the effect of the expected value of money in left MTG. The difference in lateralization of the effect of expected value between Gelskov et al. (2016) and our study can be attributed to different disorders. Paulus et al. (2005) also show activation of right MTG during the prediction phase in a two-choice prediction task in abstinent amphetamine dependent subjects. Right MTG was also seen to be involved during impulsive decision-making in a delay discounting task in smokers in Mackillop et al (2012), however, its activity was negatively correlated with the degree of discounting. A possible reason for the deactivation of middle temporal gyri during inhalation could be that these functions are not required during the inhalation phase. However, the increased connectivity between the right middle temporal gyrus and motor areas during inhalation suggests that the right MTG plays a role in drug consumption that is beyond its evaluative and predictive functions. Lawn et al. (2020) have shown greater activation of bilateral MTG during cigarette choice events as compared to rest periods in a purchase task. In relation to this, we also observe the effect of win – loss in the nicotine sessions in the right MTG. Wilson et al. (2005) show greater activation of bilateral MTG to cigarette cues as compared to neutral cues in smokers, and this difference in activation to both the cues is higher when the cigarette is available to the smokers immediately after the session as compared to when they are available 1-hour post-session. This suggests that the MTG, along with its executive functions, might also be involved in representing craving, and it deactivates during inhalation events because inhaling nicotine vapors quells the craving, at-least temporarily. Also, the inhalation event lasts for a short duration, and the subject might want to inhale as much as possible during that time, which might explain increased functional connectivity between motor regions and right MTG where previous research has shown right MTG to be involved in impulsivity (Mackillop et al., 2012).

## Limitations

There are several limitations to our study. First, the sample size in the case of the nicotine session (21) was smaller than the sample size in the case of the money session (45). With a bigger sample size and stronger control of nicotine abstinence in nicotine sessions, we might have greater power to observe the effect of gamble characteristics.

As discussed earlier, previous research has shown that reward expectancy (immediate or delayed) in an fMRI task can affect the activation in various brain regions (Wilson et al., 2005). In the current study, the nicotine reward was immediately consumable while the money reward was not. While in this study we specifically wanted to compare the effects of monetary and nicotine decision-making tasks, another study with a consumable non-drug reward such as juice or chocolates might be able to account for the effects of reward use opportunity which might have played a role in this study. Further, in case of monetary rewards, it is difficult to define what it means to ‘consume’ or ‘possess’ money. In our daily lives, we seem to assume possessing a certain amount of money when a notification on a money transfer mobile application tells us so, and thus, a message to the subjects inside the scanner that they won a certain amount of money can also be assumed to have led the subjects to believe that they possessed that money from that point onwards.

Moreover, the nicotine and money rewards presented in the task were not titrated against each other. In other words, we did not know how they compared to each other in terms of their subjective value to the subjects. If we plot the subjective value vs actual value plot for a subject, the money amounts and cigarette time duration offered to the subjects might fall at different places on the y-axis while ideally, we will want them to coincide. In future work, the reward values offered in case of drug and non-drug rewards should be matched in terms of their subjective values.

Also, the variance of the gamble outcomes and the maximum possible gamble win outcome had a strong and positive correlation (r = 0.87), implying that some of the effects of gamble variance might be mediated by the maximum possible gamble outcome. In a future study, these two gamble characteristics should be disentangled.

## Conclusion

This is the first study to look at risky decision making in smokers when the nicotine reward was available for consumption immediately after it was earned, inside the scanner. Our results suggest that using monetary rewards to study decision-making in nicotine dependence might not reflect the actual process of how individuals make the decision about using nicotine. This is because we show that risky decision-making about nicotine and monetary rewards engage different brain regions, most notably in the case of feedback where we observed regions in the cingulate cortex, and Insula representing loss over win for monetary rewards while win over loss for nicotine rewards. In the case of both the rewards, the expected value of the gambles was the best predictor of the choice behavior.

## Materials and Methods

### Participants

A total of 56 subjects were initially recruited in the study through Craigslist. The inclusion criteria were self-reported smoking of one or more packs per day and age between 18 to 40. Amongst these, 54 subjects participated in money sessions and 49 participated in nicotine sessions. During the money sessions, 2 subjects had errors during the task, 1 subject had issues with wearing the head coil, 1 subject had a tattoo that was heating up, and 5 subjects were uncomfortable in the scanner. During the nicotine sessions, 11 subjects had noisy airflow data, 11 subjects had the inhalation pipe slip during the session, 1 subject had error during the task, 1 subject had a tattoo that was heating up, and 4 subjects were uncomfortable in the scanner. After excluding the data from these subjects, we analyzed the data from a total of 46 subjects (mean = 26.2 years, SD = 6 years; 8 females) with 45 subjects corresponding to money session, and 21 subjects corresponding to nicotine session (These numbers include the 20 subjects who had data for both the money and nicotine sessions). Given the data loss and resulting difference in sample size between money and nicotine sessions, we also performed the analysis of the data from the money session of only the 20 subjects who also participated in nicotine session (supplementary material, section H).

Three subjects did not provide information about their age. All procedures were approved by the IRB of Indiana University.

### Task

The task consisted of choosing between two options: a gamble, and a sure thing, similar to a previously published task (Fukunaga et al., 2018). Figure 8 shows the information about the two options. The gamble option had three possible outcomes. The probability and the value associated with each option was explicitly shown to the subject. The sure-thing option had only one possible outcome which was shown to the subject and if this option was chosen, it resulted in the specified outcome with 100% certainty. The task presented five different gambles (table 2), such that the probability of loss, the variance of outcomes, and the expected value of the gambles were orthogonalized across the gambles (table 3). The orthogonalization of gamble properties enabled us to look at the effect of each of these properties separately. The value of the sure thing option presented with the gamble was set to be equal to the certainty equivalent (CE) for the gamble of a subject, calculated based on the subject’s choice behavior for that gamble until that instance of choice presentation. The CE is the value of the sure-thing at which the subject is indifferent between choosing the gamble and the sure-thing. If the value of the sure-thing is more than CE, the subject tends to choose sure-thing more while if it is less than CE, then the subject tends to choose the gamble more. The value of CE depends on the subject and the gamble. The process of estimating the CE values for the gambles throughout the session is the same as described in Fukunaga et al. (2018). Briefly, an initial estimate of the CE values for the gambles was obtained based on the subject’s choice behavior on the first 8 trials of each of the gambles. The functional data collection began after the initial CE for the gambles was obtained. The CE values were also adjusted throughout the session, based on the subject’s choice behavior, using a modified proportional integral derivative (PID) controller to control the probability of choosing the gamble at 50%.

**Table 2:**
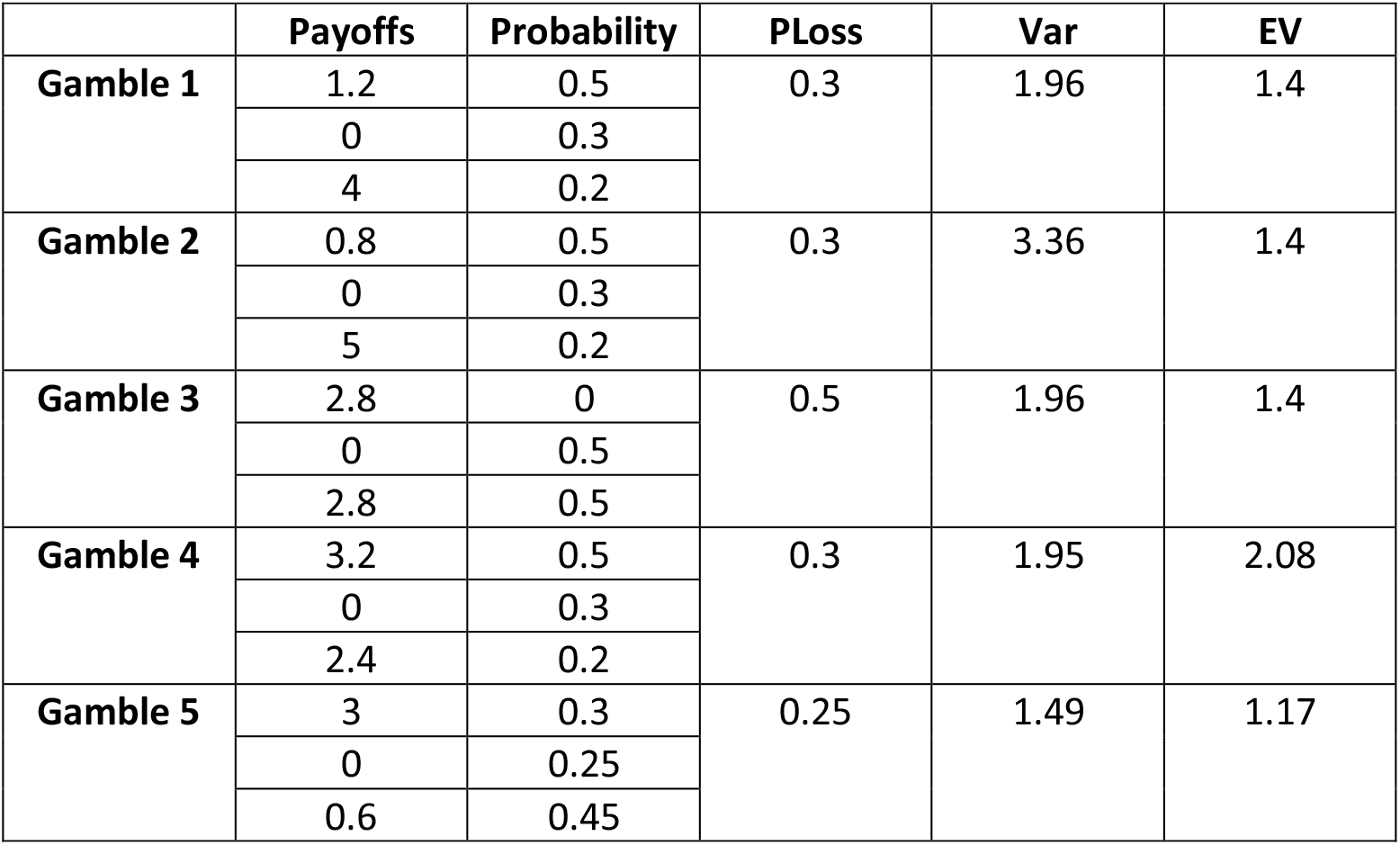
Characteristics of the five gambles; PLoss – Probability of loss, Var – Variance, EV – Expected value

|  | Payoffs | Probability | PLoss | Var | EV |
| --- | --- | --- | --- | --- | --- |
| **Gamble 1** | 1.2 | 0.5 | 0.3 | 1.96 | 1.4 |
|  | 0 | 0.3 |  |  |  |
|  | 4 | 0.2 |  |  |  |
| **Gamble 2** | 0.8 | 0.5 | 0.3 | 3.36 | 1.4 |
|  | 0 | 0.3 |  |  |  |
|  | 5 | 0.2 |  |  |  |
| **Gamble 3** | 2.8 | 0 | 0.5 | 1.96 | 1.4 |
|  | 0 | 0.5 |  |  |  |
|  | 2.8 | 0.5 |  |  |  |
| **Gamble 4** | 3.2 | 0.5 | 0.3 | 1.95 | 2.08 |
|  | 0 | 0.3 |  |  |  |
|  | 2.4 | 0.2 |  |  |  |
| **Gamble 5** | 3 | 0.3 | 0.25 | 1.49 | 1.17 |
|  | 0 | 0.25 |  |  |  |
|  | 0.6 | 0.45 |  |  |  |

**Table 3:** Correlation between gamble characteristics; PLoss – Probability of loss, Var – Variance, EV – Expected value

|  | PLoss | Var | EV |
| --- | --- | --- | --- |
| **PLoss** | - | - | - |
| **Var** | -0.016 | - | - |
| **EV** | -0.015 | 0.016 | - |

The location of the gamble and the sure thing was counterbalanced throughout the trial to avoid confounds, such that sometimes the gamble option appeared on right and sometimes it appeared on left. To indicate the choice, subjects had to press a button with the index finger of the hand corresponding to that side (For example, using the right index finger to choose the right option.)

In both the sessions, only non-negative outcomes were possible, with a non-zero outcome considered as a win and a zero-outcome considered as a loss.

In the case of the money session, the outcome was in terms of dollars. At the end of the money session, a part of the total amount paid to the subjects was proportional to the amount they won through the session (Supplementary material, section D). In the case of the nicotine session, the outcome was in terms of seconds for which subjects could inhale nicotine vapors. The subjects could inhale nicotine vapors inside the scanner in the event of winning through an MR compatible e-cigarette.

The event sequence in the task was as follows (refer to figure 8): A fixation screen, presentation of choice options, acknowledgement of the option selected by the subject by displaying a red line under the option selected for a jittered duration (lasted for 2 secs + jitter of 0, 2, 4, or 6 secs), feedback or the outcome of the selected option revealed to the subject for 1.5 seconds. In the case of the nicotine session, feedback of ‘win’ was followed by the inhalation of nicotine vapor by the subjects. After a jitter (1.5, 3.5, 5.5, or 7.5 secs), the next trial began, again starting with the presentation of both the options.

### Data collection

#### Initial breath CO

At the beginning of both money and nicotine sessions, the breath carbon monoxide (CO) of the subjects was recorded using the Smokerlyzer PiCO (Bedfont Scientific, UK), as a measure of recent smoking. This was done to find whether the subjects adhered to the guidelines for abstinence. However, none of the subjects was excluded based on initial breath CO.

#### Self-reported craving

After every 5 trials, subjects were asked to self-report their ‘desire for nicotine’ on a Likert scale of 1-10 using the buttons on the hand paddle provided to them.

### fMRI data acquisition and preprocessing

Echo Planar Imaging (EPI) was performed using the Siemens 3 Tesla TIM Trio MRI scanner at the imaging research facility at the Psychological and Brain Sciences department at Indiana University Bloomington. The functional and structural scans were collected using a 32-channel head coil, and with axial slices at an angle of 30° with the line connecting anterior commissure and posterior commissure. The latter was done to get a better signal to noise ratio. T2* weighted functional scans were collected with a TR = 2000 ms, TE = 25 ms, flip angle = 70°, and a 64 × 64 voxel matrix. For each run, 240 volumes were collected with each volume consisting of 35 slices of the thickness of 3.8 mm.

T1 weighted functional scans were collected with a TR = 1800 ms, TE = 2.7 ms, flip angle = 9°, and a 256 × 256 voxel matrix. In this case, 160 slices were collected with a thickness of 1 mm.

Subjects were scanned while they performed the task on two days, one day for the money session and one day for the nicotine session, with the days on which the money session happened, day 1 or day 2, counterbalanced across the subjects. There were 5 runs of the task, about 8 minutes each. The total number of trials was 189 on average in the case of the money session and 152 on average in the case of the nicotine session. The lesser number of trials in nicotine sessions were a result of the lengthening of the duration of nicotine trials because they involved time for inhalation whenever the subject won some reward.

Preprocessing of the functional data was performed using SPM5. Spike correction was performed using 3dDespike in AFNI, and the slice timing correction was performed in SPM5. For every subject, the functional scans were first realigned, then co-registered to the structural scan, and then normalized to the standard Montreal Neurological Institute (MNI) space. Spatial smoothing was performed using an 8 mm^3^ full-width-at-half-maximum (FWHM) kernel.

### fMRI analysis

#### 1. GLM Design

We analyzed the fMRI data using the general linear modeling (GLM) approach in SPM5. There were 12 task regressors, 24 motion regressors (6 degrees of freedom, absolute vs. differential motion for each, and the square of each resulting regressor), 1 constant for the entire session, and 1 constant for each of the five runs. The analysis was performed as a single session glm. The task regressors were: *ChoiceDur, Choice, Choice*Variance, Choice*Craving, Choice*EV, Choice*PLoss, Acknowledge, Gamble, feedbackWin, feedbackWin*Craving, feedbackLoss*, and *feedbackLoss*Craving*. We also added the estimated plasma nicotine concentration as a nuisance regressor, in the case of nicotine session, which has been described in a separate section. Moreover, a piece-wise linear interpolation was performed on the self-reported craving ratings which were recorded every five trials, to estimate the craving rating corresponding to every trial. *ChoiceDur* was an event regressor spanning the duration between option presentation and option selection. *The Choice* was an event regressor representing all the choice events, with onset time same as the time of choice, that is option selection and duration equal to zero. This was parametrically modulated by gamble characteristics and the craving rating: *Choice*Variance, Choice*Craving, Choice*EV*, and *Choice*PLoss*, entered in the same order. Here, PLoss stands for probability of loss of the gamble, Variance refers to the variance of gamble options, and EV refers to the expected value of the gamble. Similarly, event regressor *Gamble* represents all the choice events when the gamble was chosen. *Acknowledgement* represents the event of the appearance of a red line appeared under the option selected by the subject. *feedbackWin* represents event regressor representing the feedback event when the subject earned a non-zero reward with onset time as the time when the feedback was displayed to the subject and was parametrically modulated by the craving rating, leading to *feedbackWin*Craving*. Similarly, *feedbackLoss* represents the feedback event when the subject earned zero rewards and was parametrically modulated by the craving value, leading to *feedbackLoss*Craving*.

#### Obtaining plasma nicotine values – PBPK model

Blood plasma nicotine levels may exert a direct pharmacological effect on BOLD signals (Yamamoto et al., 2013), so we estimated plasma nicotine as a function of time in order to both control for it and estimate the effects of plasma nicotine on BOLD signals. The plasma nicotine values for the nicotine session were obtained by entering initial CO, nicotine vapor concentration, and nicotine vapor airflow values as a function of time into a physiologically based pharmacokinetic (PBPK) model of nicotine and cotinine in humans. We previously used a simple PBPK model to validate our device’s ability to increase blood plasma nicotine (Vélez de Mendizábal et al., 2015). For the present study, we fit a more temporally precise nine-compartment PBPK model (Robinson et al., 1992) to the previous data set (Vélez de Mendizábal et al., 2015). The PBPK model prediction of plasma nicotine in arterial blood was resampled at every fMRI volume acquisition and entered into the GLM as a plasma nicotine regressor (without convolution by a canonical hemodynamic response function).

#### Additional GLM designs

The majority of the results that are reported for the glm analysis are from the glm described above and will be referred as the **glm A** from onwards. We also formed other glms to investigate additional effects:

##### Glm B

It is the same as glm A except that it has these additional regressors to control for the motor effects of inhaling vapor: *Inhale, Inhale*Airflow*. Here *Inhale* represents the event of nicotine vapor inhalation for the duration won by the subject. Airflow is the average airflow of inhalation during a particular inhalation event. This glm was only formed for the nicotine sessions since only the nicotine session involved inhalation.

##### Glm C

It is the same as glm A except that the value of sure-thing (ST) is also added as a parametric modulator on the *Choice*, leading to an additional regressor: *Choice*ST. Choice*ST* was inserted after all the other regressors during the choice event, to allow us to estimate the effects of the ST value on BOLD activity.

It should be highlighted again that the sure-thing value presented with a gamble was the certainty equivalent value calculated for that subject based on the choice behavior on that gamble up until that particular choice event and also that there was a significant and positive correlation between sure-thing values and the expected value of the gamble. Thus, to make sure that the effect of a sure thing in these areas, was not mediated by the effect of gamble variance, probability of loss, or expected value, we had entered the sure thing regressor at the end. SPM uses a serial orthogonalization method when multiple parametrically modulated regressors are entered for a single GLM event. As a consequence, the order of parametrically modulated regressors matters – the first parametric modulator will have access to all the variance of the BOLD signal, while subsequent parametric modulators will have access only to the residual variance after that accounted for by the earlier parametric modulators is partialed out.

For all GLMs, a canonical hemodynamic response function with the order of expansion set to 1- and 16-time bins per scan was used.

#### 2. PPI analysis

Delivering the nicotine rewards inside the scanner provided us an opportunity to look at the brain regions activated when the smokers consumed nicotine vapor. These regions were determined using the glm B described above. Further, we also tested for regions that might not be directly involved in motor actions related to nicotine vapor inhalation but could be indirectly involved in nicotine consumption through increased functional connectivity with the motor regions during inhalation. We performed a Psychophysiological Interaction (PPI) analysis with the data from nicotine sessions for determining such regions (O’Reilly et al., 2012). We first formed a mask using the regions in the right motor cortex that showed a significant effect of *Inhale* regressor in glm B in the second-level analysis with all the subjects in nicotine session (We refer to it as the ‘main mask’). Then, we looked at the first level analysis of each subject to find the region within the main mask showing a significant effect of *Inhale* regressor in glm B. We performed small volume correction using the main mask to find the region corresponding to inhalation in a particular subject, and then used this region to form a mask for that subject. Simply put, for a particular subject, we found a region of overlap between the main mask and the part of the right motor cortex showing the effect of *Inhale* event regressor in that subject, and then formed a mask defining the region of this overlap. This way, we had a separate mask for each subject defining a region in the right motor cortex. We used these masks to get the preprocessed data from the voxels within the mask. This data was normalized and detrended and was used to form a ‘physiological regressor’ in SPM5. Similarly, the *Inhale* event regressor formed the ‘psychological regressor’. These both regressors were multiplied and then convolved with hemodynamic response function to get the PPI regressor. An additional GLM was then estimated in SPM5, consisting of the PPI regressor, psychological regressor, and physiological regressor, entered in that order, and 24 motion regressors. This was the PPI analysis for the right motor cortex involved in inhalation. A similar process was used to also perform the PPI analysis for the left motor cortex involved in inhalation.

Any effect in a region in the glm and PPI analyses above was considered significant, if the cluster corrected p-value of the effect was < 0.05 with the cluster defining threshold as < 0.001 and the minimum size of the cluster as at-least 5 contiguous voxels unless otherwise specified. Cluster significance was verified using a recently compiled version of 3DClustSim, to avoid alpha-inflation issues associated with earlier versions (Eklund et al., 2016).

## Behavioral analysis

The certainty equivalent of a subject for a particular gamble provides a measure of the tendency of that subject to choose the gamble and is thus a measure of risk-seeking. Note that the certainty equivalent alone cannot inform about the absolute tendency of someone to choose gamble, CE is only the value of the sure thing at which the choice probability of sure thing option and gamble option are 0.5 each. We regressed the CE values of the five gambles across all the subjects with the probability of loss, expected value, and variance of the gamble outcomes.

CE ∼ 1 + Probability of loss + Variance of gamble outcomes + Expected value

This enabled us to evaluate the gamble characteristic that most influenced the choice between gamble and sure thing option. The gamble characteristic with the greatest regression coefficient can be inferred as the one leading to a greater CE and thus, the decisions in favor of the gamble option.

Additionally, we had also done following analysis: 1. Comparing ((CE – EV)/EV) values in money session vs nicotine session (both independent and paired samples t-test); here EV refers to expected value of gambles 2. Correlation between CE values in money and nicotine sessions 3. Comparing self-reported craving values between money and nicotine sessions (both independent and paired samples t-test) 4. Comparing initial CO between money and nicotine sessions 5. Correlation between the coefficients of gamble characteristics in the regression CE ∼ 1 + Probability of loss + Variance of gamble outcomes + Expected value, with initial CO and self-reported craving. 6. Correlation between initial CO and self-reported craving of subjects across both sessions. 7. Comparing response time between money and nicotine sessions

## Author contributions

JWB, JM, LC, CP, and PF designed the study. JWB implemented the study. PM, CH, RP, EA, and LF collected and processed the data. PM and JWB analyzed the data. PM and JWB wrote the paper. All authors edited and approved the final manuscript.

## Acknowledgments

We thank J. Sturgeon for developing and building the custom nicotine vapor device and I. Innis and K. Heeter for assistance with data collection. Supported by NIH R21DA040773.

