## Supplementary material for "Neural bases of risky decisions involving nicotine vapor versus monetary reward"

#### Section A: Various task effects observed in Middle Temporal Lobe

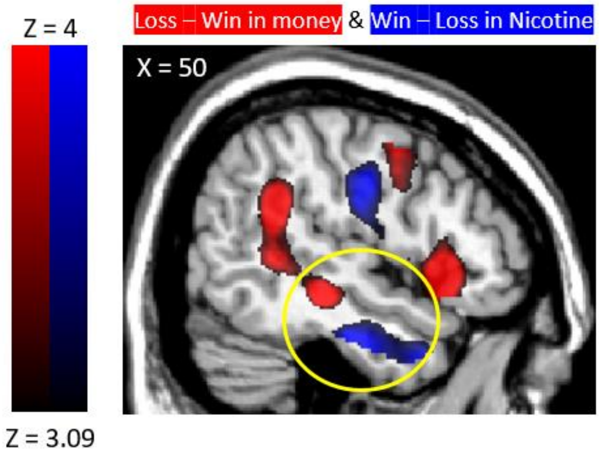

*Figure S1: Effect of loss – win in money session (red) and win – loss in nicotine session (blue) on right Middle Temporal Lobe*

Additionally, the effects of variance, also extending to cerebellum (peak voxel MNI: 22, -50, -28; -46, -56, 20), probability of loss (peak voxel MNI: 46, -44, 2; -52, -54, -4), and expected value (peak voxel MNI: 56, -38, -6; -46, -36, -12) of gambles in the money session were observed in both left and right temporal lobe. Temporal lobe also showed bilateral deactivation with nicotine inhalation. However, there seems to be a lateralization in how feedback in represented in money and nicotine sessions in temporal lobe. Right temporal lobe showed effects of loss – win in money session (peak voxel MNI: 56, -46, 22) and win – loss in nicotine session (peak voxel MNI: 6, 16, 32), while left temporal lobe showed effects of win – loss in money session (peak voxel MNI: -42, -70, 30) and loss – win in nicotine session (-66, -48, 22). This is like the error reversal effect observed in cingulate and insula, discussed in the main article.

#### Section B: Various task effects observed in motor cortex

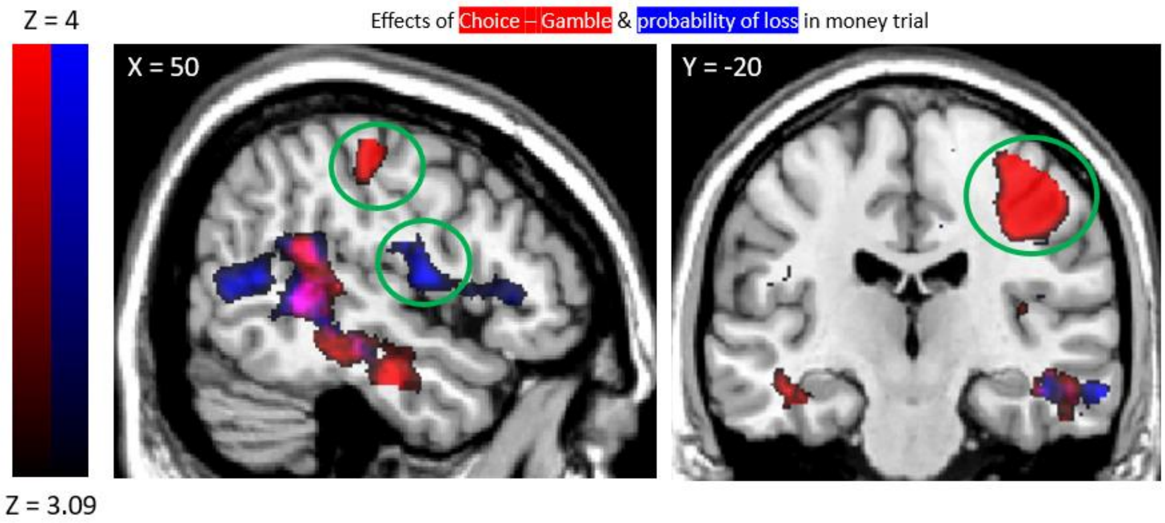

*Figure S2: Effects of Choice - Gamble contrast on motor cortex (red) (peak voxel MNI: 40, -22, 42) and positive loading on probability of loss regressor (blue) (peak voxel MNI: 52, -2, 12) on motor cortex in money session*

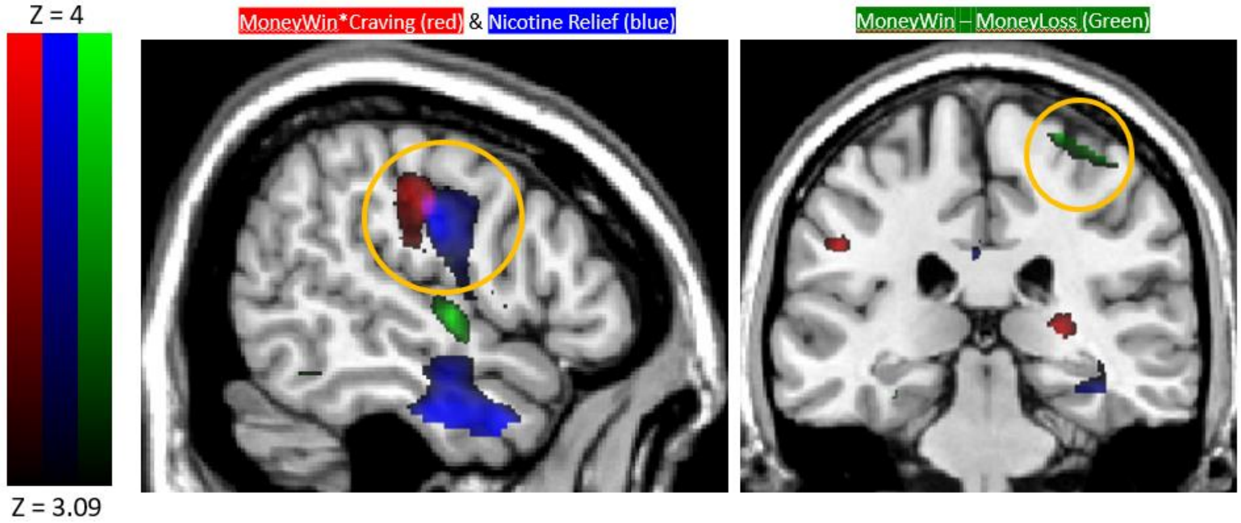

*Figure S3: Effect of Win - Loss (red) (peak voxel MNI: 26, -28, 70) and Win*Craving (blue) (peak voxel MNI: 54, -20, 38, t = 4.22, cluster size = 238, p value < 0.001, cluster corrected, two-tailed) in money session. Also, effect of relief (green) (peak voxel MNI: 46, -10, 36) in nicotine session.*

Additionally, precentral/postcentral gyri also show effect of surething value (positive effect of Choice*ST regressor; peak voxel MNI: 36, -26, 44; -50, -18, 50).

#### Section C: Results from GLM and PPI analyses

##### C.1 General Linear Modeling (GLM) analysis

### GLM A

#### *Choice – Gamble* contrast

Table S1: Positive loading on Choice – Gamble in money session

| | | | Peak MNI coordinates | | | | |
|---|---|---|---|---|---|---|---|
| **Region** | **Laterality** | **Cluster Size** | **X** | **Y** | **Z** | **Max stat t** | **P Cluster Corrected** |
| Cerebellum_4_5 left (Culmen), bilateral lingual gyrus | Bilateral | 10613 | -20 | -48 | -24 | 8.08 | <0.001 |
| | | | 16 | -72 | -6 | 7.54 | |
| Precentral/Postcentral Gyrus (BA 3) | Right | 1147 | 40 | -22 | 42 | 7.4 | <0.001 |
| Middle Frontal Gyrus/ Superior Frontal Gyrus (BA 8) | Left | 801 | -10 | 38 | 40 | 5.82 | <0.001 |
| Middle Frontal Gyrus/ Superior Frontal Gyrus (BA 9) | Right | 882 | 12 | 54 | 30 | 4.74 | <0.001 |
| Precuneus (BA 31) | Right | 215 | 14 | -50 | 34 | 4.48 | 0.012 |

Table S2: Negative loading on Choice – Gamble in money session

| | | | Peak MNI coordinates | | | | |
|---|---|---|---|---|---|---|---|
| **Region** | **Laterality** | **Cluster Size** | **X** | **Y** | **Z** | **Max stat t** | **P Cluster Corrected** |
| Precuneus (BA 7) | Bilateral | 1274 | 12 | -68 | 52 | 6.30 | <0.001 |
| | | | -18 | -74 | 42 | 5.09 | |
| Cuneus (BA 18) | Left | 285 | -10 | -100 | 2 | 5.49 | 0.003 |

Table S3: Positive loading on Choice – Gamble in nicotine session

| | | | Peak MNI coordinates | | | | |
|---|---|---|---|---|---|---|---|
| **Region** | **Laterality** | **Cluster Size** | **X** | **Y** | **Z** | **Max stat t** | **P Cluster Corrected** |
| Lingual Gyrus (BA 19) Left; small bilateral cerebellum_6 (Declive) | Bilateral | 331 | -12 | -62 | -2 | 5.48 | 0.001 |
| | | | 4 | -78 | -18 | 4.18 | |
| Posterior Cingulate (BA 30) | Right | 139 | 18 | -60 | -8 | 4.66 | 0.06 |

Table S4: Negative loading on Choice – Gamble in nicotine session

| | | | Peak MNI coordinates | | | | |
|---|---|---|---|---|---|---|---|
| **Region** | **Laterality** | **Cluster Size** | **X** | **Y** | **Z** | **Max stat t** | **P Cluster Corrected** |
| Caudate/Sub-gyral region | Right | 213 | 18 | -10 | 24 | 4.56 | 0.011 |

#### *Choice*Var* regressor

Table S5: Positive loadings on variance regressor in money session

| | | | Peak MNI coordinates | | | | |
|---|---|---|---|---|---|---|---|
| **Region (Putative)** | **Laterality** | **Cluster Size** | **X** | **Y** | **Z** | **Max stat t** | **P Cluster Corrected** |
| Bilateral Cerebellum_6 (Culmen), right Middle Temporal Gyrus | | 4726 | -28 | -50 | -28 | 6.6 | <0.001 |
| | | | 22 | -50 | -28 | 6.45 | |
| Middle/Superior Frontal Gyrus | Left | 1043 | -10 | 38 | 40 | 6.47 | <0.001 |
| Temporal lobe | Left | 157 | -46 | -36 | -12 | 5.08 | 0.042 |
| Middle/Superior Frontal Gyrus | Right | 894 | 18 | 40 | 40 | 5.03 | <0.001 |
| Cingulate Gyrus (BA 31) | Right | 172 | 10 | -42 | 32 | 4.56 | 0.03 |
| Superior Temporal Gyrus | Left | 366 | -46 | -56 | 20 | 4.51 | 0.001 |

Table S6: Negative loadings on variance regressor in money session

| | | | Peak MNI coordinates | | | | |
|---|---|---|---|---|---|---|---|
| **Region (Putative)** | **Laterality** | **Cluster Size** | **X** | **Y** | **Z** | **Max stat t** | **P Cluster Corrected** |
| Occipital Lobe (Cuneus) | Left | 452 | -12 | -98 | 2 | 5.8 | <0.001 |
| Precuneus/Superior Parietal Lobule (BA 7) | Right | 929 | 14 | -68 | 50 | 5.75 | <0.001 |
| Precuneus (BA 7) | Left | 420 | -20 | -74 | 40 | 5.07 | <0.001 |

No significant clusters for positive loading on variance regressor in nicotine session

Table S7: Negative loadings on variance regressor in nicotine session (set level p value 0.853)

| | | | Peak MNI coordinates | | | | |
|---|---|---|---|---|---|---|---|
| **Region (Putative)** | **Laterality** | **Cluster Size** | **X** | **Y** | **Z** | **Max stat t** | **P Cluster Corrected** |
| Occipital lobe (Cuneus) | Left | 165 | -18 | -94 | 16 | 4.88 | 0.025 |

#### *Choice*Ploss* regressor

Table S8: Positive loadings on probability of loss regressor in money session

| | | | Peak MNI coordinates | | | | |
|---|---|---|---|---|---|---|---|
| **Region (Putative)** | **Laterality** | **Cluster Size** | **X** | **Y** | **Z** | **Max stat t** | **P Cluster Corrected** |
| Precentral Gyrus (BA 6, Close to Insula) | Right | 857 | 52 | -2 | 12 | 5.51 | <0.001 |
| Superior/Middle Temporal Gyrus | Right | 1842 | 46 | -44 | 2 | 5.38 | <0.001 |

| | | | | | | | |
|---|---|---|---|---|---|---|---|
| (BA 22) | | | | | | | |
| Fusiform Gyrus (BA 20, BA 37), Cerebellum_4_5 (Culmen) | Left | 399 | -30 | -40 | -22 | 5.36 | <0.001 |
| Fusiform (BA 36, BA 37), Cerebellum_4_5 (Culmen) | Right | 278 | 28 | -40 | -20 | 5.2 | 0.003 |
| Middle Frontal Gyrus/ Superior Frontal Gyrus (BA 8, BA 9) | Left | 700 | -16 | 46 | 22 | 4.96 | <0.001 |
| Middle Temporal Gyrus (BA 37, BA 39), Superior Temporal Gyrus (BA 22) | Left | 1341 | -52 | -54 | -4 | 4.92 | <0.001 |
| Postcentral Gyrus (BA 40) | Right | 185 | 16 | -44 | 66 | 4.27 | 0.022 |

Table S9: Negative loadings on probability of loss regressor in money session

| | | | **Peak MNI coordinates** | | | | |
|---|---|---|---|---|---|---|---|
| **Region (Putative)** | **Laterality** | **Cluster Size** | **X** | **Y** | **Z** | **Max stat t** | **P Cluster Corrected** |
| Cuneus, Middle Occipital Gyrus (BA 17, BA 18), Lingual Gyrus | Bilateral | 2052 | 14 | -96 | 6 | 8.45 | <0.001 |
| | | | -18 | -88 | -12 | 5.56 | |
| Superior Parietal Lobule/Precuneus (BA 7) | | 189 | -10 | -72 | 54 | 4.84 | 0.02 |
| Superior Parietal Lobule/Precuneus (BA 7) | | 307 | 20 | -68 | 50 | 4.74 | 0.002 |

No significant clusters for positive loading on probability of loss regressor in nicotine session

Table S10: Negative loadings on **probability of loss** regressor in nicotine session (set level p value is ~ 0.5)

| | | | **Peak MNI coordinates** | | | | |
|---|---|---|---|---|---|---|---|
| **Region (Putative)** | **Laterality** | **Cluster Size** | **X** | **Y** | **Z** | **Max stat t** | **P Cluster Corrected** |
| Cingulate Gyrus right; Caudate left, Corpus Callosum left | Bilateral | 11371 | 18 | -14 | 28 | 8.98 | <0.001 |
| | | | -8 | -10 | 24 | 7.83 | |
| Brodmann Area 18 (Occipital Lobe; Cuneus) | Right | 393 | 22 | -90 | 16 | 5.64 | <0.001 |

***Choice*EV regressor*** **regressor**

Table S11: Positive loadings on expected value regressor in money session

| | | | Peak MNI coordinates | | | | |
|---|---|---|---|---|---|---|---|
| **Region (Putative)** | **Laterality** | **Cluster Size** | **X** | **Y** | **Z** | **Max stat t** | **P Cluster Corrected** |
| Cerebellum_6 (Culmen) | Bilateral | 1778 | -26 | -48 | -26 | 6.89 | <0.001 |
| | | | 26 | -48 | -26 | 6.04 | |
| Middle Temporal Lobe | Left | 411 | -46 | -36 | -12 | 5.59 | <0.001 |
| Middle Temporal Gyrus | Right | 182 | 56 | -38 | -6 | 5.21 | 0.027 |

Table S12: Negative loadings on expected value regressor in money session

| | | | Peak MNI coordinates | | | | |
|---|---|---|---|---|---|---|---|
| **Region (Putative)** | **Laterality** | **Cluster Size** | **X** | **Y** | **Z** | **Max stat t** | **P Cluster Corrected** |
| Superior Parietal Lobule (Brodmann Area 7) | Right | 553 | 22 | -68 | 56 | 5.17 | <0.001 |
| Cuneus | Left | 180 | -10 | -98 | 2 | 4.94 | 0.028 |

No significant clusters for positive loading on expected value regressor in nicotine session (a cluster in OFC/vmPFC was visible but it did not pass cluster correction)

No significant clusters for negative loading on expected value regressor in nicotine session

##### *Choice*Craving* regressor

No significant clusters for positive loading on craving regressor in money session.

Table S13: Negative loadings on craving regressor in money session

| | | | Peak MNI coordinates | | | | |
|---|---|---|---|---|---|---|---|
| **Region (Putative)** | **Laterality** | **Cluster Size** | **X** | **Y** | **Z** | **Max stat t** | **P Cluster Corrected** |
| Lingual Gyrus | Right | 196 | 22 | -62 | -10 | 5.68 | 0.006 |
| Superior Parietal Lobule (BA 40) | Right | 126 | 36 | -56 | 56 | 5.33 | 0.043 |

No significant clusters for positive or negative loading on craving regressor in nicotine session.

We also replaced interpolated self-reported craving as a measure of craving with deduced craving in case of nicotine session. Deduced craving was an indicator of proportion of inhalation duration won by a subject, in which the subject inhaled the vapors. Deduced craving obtained from inhalation of one session was taken as measure of craving during the next session. We did not find any significant clusters for positive or negative loading on craving regressor in nicotine session in this case as well.

##### *FeedbackLoss – FeedbackWin* contrast

Table S14: Positive loading on feedbackLoss – feedbackWin in money session

| | | | Peak MNI coordinates | | | | |
|---|---|---|---|---|---|---|---|
| **Region (Putative)** | **Laterality** | **Cluster Size** | **X** | **Y** | **Z** | **Max stat t** | **P Cluster Corrected** |
| Insula, Inferior Frontal Gyrus (BA 13, BA 45, BA 47), Middle Frontal Gyrus (BA 9), Precentral Gyrus (BA 6) | Right | 2584 | 42 | 16 | 0 | 8.4 | <0.001 |
| Insula, Inferior Frontal Gyrus (BA 13, BA 47) | Left | 1234 | -32 | 16 | -10 | 7.86 | <0.001 |
| Supramarginal Gyrus, Middle Temporal Gyrus | Right | 1360 | 56 | -46 | 22 | 7.51 | <0.001 |
| Superior Frontal Gyrus/Middle Frontal Gyrus | Right | 658 | 28 | 56 | 16 | 7.42 | <0.001 |
| Superior Frontal Gyrus, Middle Frontal Gyrus (BA 8, BA 9) Cingulate Gyrus (BA 32) | Bilateral | 3409 | 14<br>-8 | 8<br>0 | 62<br>66 | 7.16<br>5.13 | <0.001 |
| Supramarginal Gyrus | Left | 378 | -58 | -48 | 28 | 6.8 | 0.001 |
| Precuneus (BA 7) | Left | 597 | -28 | -54 | 48 | 6.34 | <0.001 |
| Precuneus (BA 7) | Right | 267 | 14 | -66 | 40 | 6.13 | 0.006 |
| Cuneus (BA 18), Middle Occipital Gyrus | Left | 505 | -10 | -102 | 2 | 5.79 | <0.001 |

Table S15: Negative loading on feedbackLoss – feedbackWin in money session (This is positive loading on feedbackWin – feedbackLoss in money session)

| | | | Peak MNI coordinates | | | | |
|---|---|---|---|---|---|---|---|
| **Region (Putative)** | **Laterality** | **Cluster Size** | **X** | **Y** | **Z** | **Max stat t** | **P Cluster Corrected** |
| Posterior Cingulate, Lingual Gyrus, Cerebellum_4_5_left (Culmen) | Bilateral | 3303 | -8<br>8 | -56<br>-52 | 8<br>12 | 7.68<br>7.18 | <0.001 |
| Middle Temporal Gyrus/Angular Gyrus (BA 39) | Left | 367 | -42 | -70 | 30 | 7.11 | 0.001 |
| Superior Temporal Gyrus (BA 22) | Right | 493 | 62 | -8 | 0 | 6.23 | <0.001 |
| Middle Frontal Gyrus (BA 8) | Left | 561 | -22 | 18 | 52 | 5.89 | <0.001 |
| Middle Temporal Gyrus (BA 21), Superior Temporal Gyrus (BA 22) | Left | 165 | -58 | -8 | -4 | 4.57 | 0.045 |
| Middle Frontal Gyrus (BA 10, BA 11) | Bilateral | 294 | 0<br>-2 | 42<br>46 | -14<br>-12 | 4.33<br>4.30 | 0.004 |
| Precentral Gyrus (BA 4) | Right | 167 | 26 | -28 | 70 | 4.11 | 0.044 |

Table S16: Positive loading on feedbackLoss – feedbackWin in nicotine session

| | | | Peak MNI coordinates | | | | |
|---|---|---|---|---|---|---|---|
| **Region (Putative)** | **Laterality** | **Cluster Size** | **X** | **Y** | **Z** | **Max stat t** | **P Cluster Corrected** |
| Superior Temporal Gyrus/Supramarginal Gyrus | Left | 324 | -66 | -48 | 22 | 6.19 | 0.027 |

Table S17: Negative loading on feedbackLoss – feedbackWin in nicotine session (This is positive loading on feedbackWin – feedbackLoss in nicotine session)

| | | | Peak MNI coordinates | | | | |
|---|---|---|---|---|---|---|---|
| **Region (Putative)** | **Laterality** | **Cluster Size** | **X** | **Y** | **Z** | **Max stat t** | **P Cluster Corrected** |
| Insula/Cingulate Gyrus, right Temporal pole/Right Middle Temporal Gyrus | Bilateral | 11934 | -36 | -6 | 12 | 7.16 | <0.001 |
| | | | 6 | 16 | 32 | 6.68 | |
| Cerebellum_6 (Declive) | Bilateral | 332 | 14 | -66 | -22 | 5.78 | 0.025 |
| | | | -12 | -62 | -24 | 4.65 | |

#### (loss – win)*craving contrast

No significant clusters for positive and negative loading on (loss – win)*craving contrast in money session.

Table S18: Positive loadings on (loss-win)*craving contrast in nicotine session

| | | | Peak MNI coordinates | | | | |
|---|---|---|---|---|---|---|---|
| **Region (Putative)** | **Laterality** | **Cluster Size** | **X** | **Y** | **Z** | **Max stat t** | **P Cluster Corrected** |
| Anterior Cingulate (BA 32) | Bilateral | 533 | 4 | 28 | 28 | 5.05 | <0.001 |

No significant clusters for negative loading on (loss – win)*craving in money session.

#### *Ngml* regressor in nicotine session

No significant clusters for positive loading on Ngml (blood plasma nicotine) regressor in nicotine session

Table S19: Negative loadings on Ngml regressor (blood plasma nicotine) in nicotine session

| | | | Peak MNI coordinates | | | | |
|---|---|---|---|---|---|---|---|
| **Region (Putative)** | **Laterality** | **Cluster Size** | **X** | **Y** | **Z** | **Max stat t** | **P Cluster Corrected** |
| Occipital Lobe | Left | 352 | -26 | -76 | 22 | 4.32 | 0.013 |

### GLM B

#### *Inhale* regressor

Table S20: Positive loading on Inhale event regressor in nicotine session

| | | | Peak MNI coordinates | | | | |
|---|---|---|---|---|---|---|---|
| **Region (Putative)** | **Laterality** | **Cluster Size** | **X** | **Y** | **Z** | **Max stat t** | **P Cluster Corrected** |
| Cerebellum (Culmen/Declive) | Left | 518 | -12 | -68 | -16 | 6.03 | 0.001 |
| Right Motor cortex | Right | 4980 | 68 | -8 | 12 | 10.23 | <0.001 |
| Left Motor cortex | Left | 3276 | -64 | -12 | 14 | 8.25 | <0.001 |

Table S21: Negative loading on Inhale event regressor in nicotine session

| | | | Peak MNI coordinates | | | | |
|---|---|---|---|---|---|---|---|
| **Region (Putative)** | **Laterality** | **Cluster Size** | **X** | **Y** | **Z** | **Max stat t** | **P Cluster Corrected** |
| Precentral, Postcentral Gyrus/MFG/IFG/ Anterior Cingulate (BA 2, 4, 32, 47, 8, 9) | Bilateral | 15176 | 46 | -30 | 50 | 9.19 | <0.001 |
| | | | -36 | -36 | 42 | | |
| Middle Temporal Gyrus | Left | 902 | -58 | -28 | -6 | 5.93 | <0.001 |
| Middle Temporal Gyrus | Right | 210 | 54 | -30 | -10 | 5.17 | 0.05 |

### GLM C:

Choice*ST regressor

Table S22: Positive loading on sure thing regressor in money session

| | | | Peak MNI coordinates | | | | |
|---|---|---|---|---|---|---|---|
| **Region (Putative)** | **Laterality** | **Cluster Size** | **X** | **Y** | **Z** | **Max stat t** | **P Cluster Corrected** |
| Cingulate Gyrus, Anterior Cingulate (BA 24 left, BA 32), Middle Frontal Gyrus (BA 9) | Bilateral | 457 | -6 | 32 | 32 | 6.39 | <0.001 |
| | | | 4 | 36 | 24 | 5.05 | |
| Cingulate Gyrus (BA 23, BA 31) | Bilateral | 437 | -4 | -36 | 34 | 6.00 | <0.001 |
| | | | 6 | -32 | 30 | 4.91 | |
| Precuneus (BA 7), Cingulate Gyrus | Bilateral | 366 | 8 | -60 | 32 | 4.69 | <0.001 |
| | | | -8 | -66 | 32 | 4.17 | |
| Postcentral gyrus (BA 3), Precentral gyrus (BA 4) | Right | 369 | 36 | -26 | 44 | 4.68 | <0.001 |

Table S23: Negative loading on sure thing regressor in money session

| | | | Peak MNI coordinates | | | | |
|---|---|---|---|---|---|---|---|
| **Region (Putative)** | **Laterality** | **Cluster Size** | **X** | **Y** | **Z** | **Max stat t** | **P Cluster Corrected** |
| Postcentral gyrus (BA 2, BA 3) | Left | 220 | -50 | -18 | 50 | 4.91 | 0.009 |

No significant clusters for both, positive and negative loading on sure thing regressor in nicotine session

#### GLM D: Same as glm A, except that the feedbackWin and feedbackLoss regressors were combined into a single feedback event regressor. In addition to craving, it was also modulated by regret. Regret = sure thin value – gamble outcome, for sessions when gamble was chosen; Regret = 0, for sessions when sure thing was chosen.

##### *Feedback*regret* regressor

No significant clusters for positive loading on feedback*regret regressor in money session

Table S24: Positive loading on regret regressor money session

| | | | Peak MNI coordinates | | | | |
|---|---|---|---|---|---|---|---|
| **Region (Putative)** | **Laterality** | **Cluster Size** | **X** | **Y** | **Z** | **Max stat t** | **P Cluster Corrected** |
| Superior Frontal Gyrus, Middle Frontal Gyrus (BA 6) | Right | 714 | 12 | 20 | 52 | 5.85 | <0.001 |

Table S25: Negative loading on feedback*regret regressor in money session (Represents of relief in money session)

| | | | Peak MNI coordinates | | | | |
|---|---|---|---|---|---|---|---|
| **Region (Putative)** | **Laterality** | **Cluster Size** | **X** | **Y** | **Z** | **Max stat t** | **P Cluster Corrected** |
| Posterior Cingulate**,** Precuneus (BA 23, BA 30, BA 31) | Bilateral | 1048 | 8 | -50 | 24 | 6.11 | <0.001 |
| | | | -6 | -52 | 18 | 5.64 | |
| Middle Occipital Gyrus (BA 18) | Right | 450 | 38 | -90 | 0 | 4.86 | <0.001 |
| Caudate | Right | 166 | 10 | 2 | 0 | 4.86 | 0.042 |
| Corpus, Callosum, Anterior Cingulate (BA 24, BA 32) | Bilateral | 368 | -2 | 30 | 8 | 4.83 | 0.001 |
| | | | 4 | 42 | 16 | 3.67 | |
| Inferior Occipital Gyrus, Fusiform gyrus (BA 18) | Left | 165 | -22 | -90 | -18 | 4.17 | 0.043 |

Table S26: Positive loadings on feedback*regret regressor in nicotine session

| | | | Peak MNI coordinates | | | | |
|---|---|---|---|---|---|---|---|
| **Region (Putative)** | **Laterality** | **Cluster Size** | **X** | **Y** | **Z** | **Max stat t** | **P Cluster Corrected** |
| Superior Parietal Lobule | Right | 217 | 32 | -72 | 44 | 5.67 | 0.049 |

Table S27: Negative loadings on feedback*regret regressor in nicotine session (Represents relief in nicotine session)

| | | | Peak MNI coordinates | | | | |
|---|---|---|---|---|---|---|---|
| **Region (Putative)** | **Laterality** | **Cluster Size** | **X** | **Y** | **Z** | **Max stat t** | **P Cluster Corrected** |
| Middle Temporal Gyrus/Inferior Temporal Gyrus | Right | 3535 | 46 | -10 | -30 | 6.78 | <0.001 |
| Insula | | | 38 | 6 | 10 | 5.88 | |
| Precentral Gyrus | | | 46 | -10 | 36 | 5.66 | |
| Cerebellum_6 (Declive) | Bilateral | 208 | 16 | -64 | -26 | 6.38 | 0.056 |
| | | | -12 | -64 | -24 | 4.28 | |

GLM E: Same as glm C, except that here, gamble choice events and sure-thing choice events were modeled separately for money session, and were each parametrically modulated by value of sure thing, leading to three additional regressors: *Gamble*ST*, *Surething*, and *Surething*ST*

Table S28: Positive loadings on contrast *Gamble*ST – Surething*ST* in money session

| | | | Peak MNI coordinates | | | | |
|---|---|---|---|---|---|---|---|
| **Region (Putative)** | **Laterality** | **Cluster Size** | **X** | **Y** | **Z** | **Max stat t** | **P Cluster Corrected** |
| Cuneus (BA 18) | Left | 798 | -12 | -102 | 4 | 5.29 | <0.001 |
| Middle Occipital Gyrus (BA 18) | Right | 373 | 36 | -92 | -4 | 4.57 | 0.001 |

No significant clusters showed negative loading on contrast *Gamble*ST – Surething*ST* in money session

#### C.2 Effects in the ‘motivated’ sub-group of subjects in nicotine session

##### ***Choice*PLoss*** **regressor**

Table S29: Positive loading on probability of loss regressor in nicotine session

| | | | Peak MNI coordinates | | | | |
|---|---|---|---|---|---|---|---|
| **Region (Putative)** | **Laterality** | **Cluster Size** | **X** | **Y** | **Z** | **Max stat t** | **P Cluster Corrected** |
| Parietal Lobe (BA 7) | Left | 169 | -22 | -78 | 40 | 5.98 | 0.002 |

Table S30: Negative loading on probability of loss regressor in nicotine session

| | | | Peak MNI coordinates |
|---|---|---|---|

| Region (Putative) | Laterality | Cluster Size | X | Y | Z | Max stat t | P Cluster Corrected |
|---|---|---|---|---|---|---|---|
| Caudate | Bilateral | 5054 | -12 | -4 | 24 | 10.68 | <0.001 |
| | | | 16 | -11 | 30 | | |
| Middle Occipital Gyrus (BA 19) | Left | 2642 | -26 | -82 | 14 | 9.33 | <0.001 |
| Middle Occipital Gyrus | Right | 542 | 26 | -80 | 14 | 8.04 | <0.001 |

##### ***FeedbackLoss – FeedbackWin* contrast**

Table S31: Positive loading on feedbackloss – feedbackwin contrast in nicotine session

| | | | Peak MNI coordinates | | | | |
|---|---|---|---|---|---|---|---|
| **Region (Putative)** | **Laterality** | **Cluster Size** | **X** | **Y** | **Z** | **Max stat t** | **P Cluster Corrected** |
| Superior Occipital Gyrus (A small tip of MTL in far back, BA 19) | Right | 1155 | 34 | -82 | 32 | 7.82 | <0.001 |

Table S32: Negative loading on feedbackloss – feedbackwin contrast in nicotine session (This is positive loading on feedbackWin – feedbackLoss in nicotine session).

| | | | Peak MNI coordinates | | | | |
|---|---|---|---|---|---|---|---|
| **Region (Putative)** | **Laterality** | **Cluster Size** | **X** | **Y** | **Z** | **Max stat t** | **P Cluster Corrected** |
| Precentral gyrus, Postcentral gyrus, BA 6, BA 43, IFG, Insula | Left | 3634 | -52 | -4 | 34 | 9.61 | <0.001 |
| Inferior Temporal Gyrus/Middle Temporal Gyrus | Right | 232 | 48 | 12 | -34 | 6.39 | 0.042 |
| Cingulate Gyrus, BA 24 | Bilateral | 590 | 2 | -8 | 32 | 5.59 | <0.001 |
| | | | -8 | 12 | 34 | | |

#### C.3 PPI analysis

Table S33: Positive loading on right PPI in nicotine session

| | | | Peak MNI coordinates | | | | |
|---|---|---|---|---|---|---|---|
| **Region (Putative)** | **Laterality** | **Cluster Size** | **X** | **Y** | **Z** | **Max stat t** | **P Cluster Corrected** |
| Middle Temporal Gyrus | Right | 375 | 52 | -42 | -2 | 5.07 | <0.001 |

No significant clusters for positive loading on left PPI in nicotine session

Table S34: Negative loading on right PPI in nicotine session

| | | | Peak MNI coordinates | | | | |
|---|---|---|---|---|---|---|---|
| **Region (Putative)** | **Laterality** | **Cluster Size** | **X** | **Y** | **Z** | **Max stat t** | **P Cluster Corrected** |
| Precentral Gyrus | Left | 549 | -58 | -6 | 26 | 5.04 | <0.001 |
| Precentral Gyrus | Right | 208 | 58 | -4 | 28 | 4.46 | 0.012 |

Table S35: Negative loading on left PPI in nicotine session

| | | | Peak MNI coordinates | | | | |
|---|---|---|---|---|---|---|---|
| **Region (Putative)** | **Laterality** | **Cluster Size** | **X** | **Y** | **Z** | **Max stat t** | **P Cluster Corrected** |
| Postcentral Gyrus | Left | 196 | -58 | -12 | 26 | 5.96 | 0.014 |
| Precentral Gyrus | Right | 178 | 62 | -12 | 32 | 4.93 | 0.021 |

#### Section D: Calculation of payment amount to the subjects, in money session

Subjects were paid $25/hr. There were also following bonuses:

- on-time bonus of $10
- $20 – (amount of CO blown into breathanalyzer, in ppm); this was a bonus was abstaining from smoking

Additionally, they also received an amount proportional to the rewards the earned during money session: Every run could lead to a maximum of 5$; an average of amount for all runs was calculated and doubled. This was the amount they received.

#### Section E: Characterization of motivated vs. unmotivated subjects.

To further ascertain the validity of the assumption that subjects wanted the rewards, we also looked at the CE trajectories of individual subjects in both money and nicotine. This revealed a sub-group in nicotine sessions in which the CE values either declined over the sessions and eventually became zero or increased over initial sessions and remained constant at a certain value. The former was the case when subjects chose the sure thing more often than the gamble and the latter was the case when subjects chose the gamble more often than the sure thing. It is hard to conclude whether the subjects in this sub-group wanted nicotine, as their behavior was not consistent with maximizing reward. We refer to these as the 'unmotivated' subjects in the nicotine session. The mean initial CO expired by the subjects (presumably indicating their satiety before the experiment) in this sub-group was more than the mean initial CO expired by other subjects in nicotine session, however, the difference is not significant. The difference between the mean self-reported craving of the sub-group with unusual choice behavior and other subjects in nicotine session is significant, with the sub-group with unusual choice behavior having less self-reported craving ($t = -3$, $p = 0.007$). This supports the possibility that the greater satiety (indicated by high initial CO) before the experiment led to unusual choice behavior. The regression of CE values with gamble characteristics for the sub-group with unusual choice behavior in nicotine session revealed a positive but weaker and non-significant effect of the expected value of gamble, as shown in table 2. Surprisingly, the effect of gamble variance in this sub-group was strong, positive, and significant (b = 2.04, p = 0.05) indicating a greater likelihood of choosing the gamble when gamble variance was high. The results from glm analyses of the 'motivated' subjects can be found in section C.2.

Table S16 Regression coefficients of probability of loss, variance and expected value of gambles for the certainty equivalent values of the gambles in unmotivated subgroup in nicotine session

| | Beta value | P value |
|---|---|---|
| **Constant** | 0.856 | 0.568 |
| **PLoss** | -2.315 | 0.340 |
| **Var** | 2.036* | 0.046 |
| **EV** | 0.302 | 0.659 |

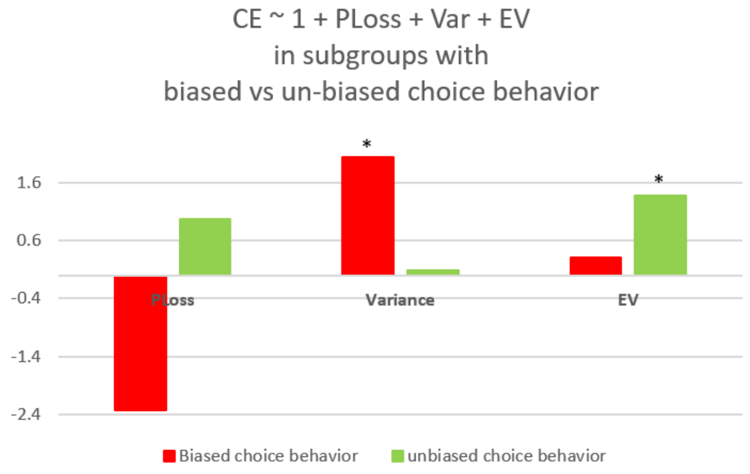

*Figure S4 Comparing effects of gamble characteristics on certainty equivalent of gambles between unmotivated subgroup (choice probability != 0.5) and motivated subgroup (choice probability = 0.5).*

#### Section F: Additional behavioral analyses

We investigated whether the effects of gamble preferences across different subjects varied depending on their initial CO, indicating their initial satiety. These provide measures of risk and reward seeking preferences as a function of nicotine satiety. The correlation between the beta weights on expected value for individual subjects and their initial CO is negative for nicotine sessions but is not significant. This correlation with EV in the case of money sessions, is negative and significant (r = -0.348, p = 0.019). The correlation between beta weights for gamble variance of individual subjects and their initial CO is positive but is not significant in the case of nicotine sessions. This correlation with gamble variance is however positive and significant (r = 0.42, p = 0.004) in the case of money sessions. We also performed a regression of CE values with the initial CO of subjects as an additional regressor which revealed a positive and significant (Nicotine: b = 0.06, p < 0.001; Money: b= 0.01, p < 0.001) effect indicating a greater likelihood of choosing the gamble (or greater risk-seeking) in subjects with greater consumption of nicotine before the scanning.

**Other behavioral analyses**: There was a positive and significant correlation between CE values in money and nicotine sessions (r = 0.526, p = 0.017). Self-reported craving of subjects in the money session was significantly more than the self-reported craving of subjects in the nicotine session (t = 3.9, p = 0.0002). This is as expected, because the subjects in the nicotine session received nicotine, which should reduce craving. Initial CO was more in the case of the money session, but the difference was not significant. The response time was more in money and significantly so when we performed paired samples t-test (t= 2.88, p = 0.016). Other behavioral analyses which were described in methods section were non-significant. The significance of all behavioral analyses was assessed with an alpha value of 0.05, not corrected for multiple comparisons.

#### Section G: Discussion of cerebellar effects

#### Effects in the cerebellum

While we did not have strong prior hypotheses regarding cerebellar involvement in addiction, several task effects were observed there during both money and nicotine sessions. During the money session, the cerebellum showed effects of the choice minus gamble contrast and of gamble characteristics -- expected value, probability of loss, and variance of gamble outcomes, suggesting a role in option evaluation and choice behavior in case of monetary rewards. During the nicotine session, the cerebellum showed effects of the inhalation event and win minus loss contrast, suggesting a role in outcome evaluation in the case of nicotine rewards. The peak MNI coordinates showing the effects of the variance and the choice minus gamble contrast in the money session are close. The choice minus gamble contrast essentially represents the effect of choosing the sure-thing option. These effects suggest that the cerebellum could be involved in driving the choice behavior away from risky options, specifically the options with a greater variance of outcomes. Cservenka and Nagel (2012)also report cerebellar activity corresponding to risky decision making in the wheel of fortune task, in a study aimed at understanding differences in risk-seeking between adolescents with a family history of alcoholism and controls. Moreover, Krain et al. (2006) in a meta-analysis of several studies of decision-making, report regions in bilateral cerebellum involved in representing risk, and implicated in decision-making in general.

The cerebellar region that showed activation during nicotine inhalation is close to the cerebellar region that was shown to be active during naturalistic smoking in Wall et al. (2017). Zubieta et al. (2005), in a PET study, also showed an increase in rCBF in the cerebellum after smoking the first cigarette, post 12 hours of abstinence in smokers.

Gelskov et al. (2016) compared the performance of pathological gamblers and healthy controls on a gambling task where they had to accept or reject a gamble with a 0.5 probability of gain and loss each. The ratio of gain and loss was varied across different gambles. They found that clusters in both anterior and posterior cerebellum showed greater activation with a greater gain/loss ratio of the gamble. This is consistent with our finding of positive loading on the expected value of gamble in the cerebellum. Moreover, Rose et al. (2013) also found the effect of the magnitude of monetary reward indicated by reward predicting stimuli in the cerebellum in smokers. They also found greater activation in the cerebellum for successful trials as compared to unsuccessful trials during the outcome phase in the MID task. The successful minus unsuccessful contrast in Rose et al. (2013) seems similar to the win minus loss contrast here, however, in our study we did not find the effect of this contrast in the case of monetary rewards, and only found it in the case of nicotine rewards. The fact that the successful trials in Rose et al. (2013) also include trials in which money was lost, suggests that the successful minus unsuccessful contrast in Rose et al. (2013) and the win minus loss contrast in the current study might not be directly comparable.

The cerebellum is generally considered as a region predominantly involved in motor control; however, several studies, suggest its roles in non-motor and cognitive functions, like emotional processing, language processing, spatial processing, working memory, and executive function (Ernst et al., 2002; Moulton et al., 2014; Stoodley, 2012). Our observations of effects in cerebellum suggest its role in decision making, not only in the case of monetary rewards but also nicotine rewards. Specifically, based on our results, the cerebellum seems to be engaged in option evaluation in case of monetary rewards while in outcome evaluation in the case of nicotine rewards. However, it is possible that with a larger sample size in the case of nicotine sessions, we might be able to observe the effects of option evaluation in the case of nicotine rewards as well. Moreover, it is also noteworthy that we did not observe any effects in the outcome phase in the case of monetary rewards in the cerebellum while only observed them in the case of nicotine rewards. Also, the previous studies with monetary rewards that were discussed in this section also did not report cerebellar activation during the outcome phase while the study involving cigarettes did. One possible explanation for this can be found in studies of nicotine dependence in rodents. The molecular biology studies in mice have shown that the activation of the Medial Habenula-Interpeduncular Nucleus pathway is involved in alleviating withdrawal symptoms upon nicotine consumption (McLaughlin et al., 2017). Extrapolating these findings to nicotine dependence in humans suggests that we would expect at-least the activation of cerebellar peduncles to be exclusively observed during nicotine win and inhalation (and not money win).

#### Section H: Money session results for subjects that had both the money and nicotine sessions.

In the current study, there are many more subjects in money session (45) than nicotine session (21). In this section we report the results that were discussed in the main article for money session but with only 20 subjects that participated in both money and nicotine sessions. This would provide a fairer comparison with results from nicotine session. With only 20 subjects, we observe that the effect of variance during choice is not as strong in Middle/Superior Frontal gyrus. Many other effects become weaker too, as expected. However, we still observe the error reversal effect even with this smaller set of subjects, as shown in figure S5.

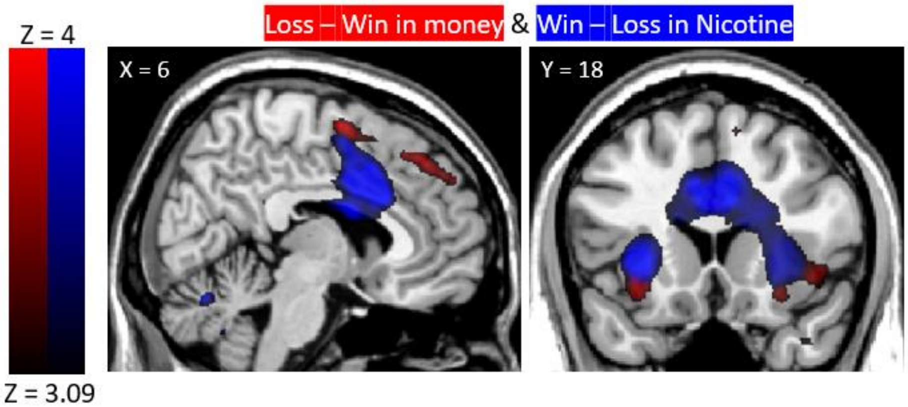

*Figure S5: Feedback effect reversal in money vs nicotine session when money session analysis was performed only for 20 subjects having both money and nicotine session data. Regions that load significantly more on money loss than money win are shown in re*

#### GLM A:

##### *Choice – Gamble* contrast

Table S37: Positive loading on Choice – Gamble in money session (n = 20)

| | | | Peak MNI coordinates | | | | |
|---|---|---|---|---|---|---|---|
| **Region** | **Laterality** | **Cluster Size** | **X** | **Y** | **Z** | **Max stat t** | **P Cluster Corrected** |
| Cerebellum_6, Cerebellum_4_5 (Culmen), Fusiform gyrus | Left | 716 | -24 | -54 | -24 | 7.32 | <0.001 |
| Precentral/Postcentral Gyrus (BA 3, BA 6) | Right | 552 | 38 | -12 | 56 | 7.19 | <0.001 |
| Fusiform gyrus | Right | 110 | 34 | -42 | -24 | 5.92 | 0.09 |
| Occipital lobe, Lingual Gyrus (BA 18) | Left | 124 | -8 | -78 | -8 | 5.06 | 0.06 |

##### *Choice*Var* regressor

Table S38: Positive loadings on variance regressor in money session (n = 20)

| | | | Peak MNI coordinates | | | | |
|---|---|---|---|---|---|---|---|
| **Region (Putative)** | **Laterality** | **Cluster Size** | **X** | **Y** | **Z** | **Max stat t** | **P Cluster Corrected** |
| Bilateral Cerebellum_4_5 (Culmen), Fusiform gyrus | Right | 190 | 34 | -42 | -24 | 6.17 | 0.009 |
| Cerebellum_6, Cerebellum_4_5 | Left | 378 | -26 | -54 | -26 | 6.05 | <0.001 |
| Temporal lobe | Left | 142 | -46 | -36 | -16 | 5.81 | 0.033 |

##### *Choice*Ploss* regressor

Table S39: Positive loadings on probability of loss regressor in money session (n = 20)

| | | | Peak MNI coordinates | | | | |
|---|---|---|---|---|---|---|---|
| **Region (Putative)** | **Laterality** | **Cluster Size** | **X** | **Y** | **Z** | **Max stat t** | **P Cluster Corrected** |
| Middle Frontal Gyrus/ Superior Frontal Gyrus (BA 8, BA 9) | Left | 639 | -20 | 42 | 16 | 6.29 | <0.001 |
| Insula (BA 13) | Left | 654 | -38 | 4 | 12 | 5.77 | <0.001 |
| Cerebellum_4_5 (Culmen) | Left | 263 | -8 | -56 | -8 | 5.62 | 0.001 |
| Inferior Parietal Lobule, Supramarginal gyrus | Left | 124 | -42 | -40 | 28 | 4.8 | 0.048 |

#### ***Choice*EV regressor* regressor**

Table S40: Positive loadings on expected value regressor in money session (n = 20)

| | | | Peak MNI coordinates | | | | |
|---|---|---|---|---|---|---|---|
| **Region (Putative)** | **Laterality** | **Cluster Size** | **X** | **Y** | **Z** | **Max stat t** | **P Cluster Corrected** |
| Cerebellum_6 (Culmen) | Left | 143 | -30 | -48 | -30 | 6.22 | 0.037 |
| Temporal Lobe (BA 37) | Left | 112 | -46 | -46 | -14 | 5.55 | 0.088 |

#### ***FeedbackLoss – FeedbackWin* contrast**

Table S41: Positive loading on feedbackLoss – feedbackWin in money session (n = 20)

| | | | Peak MNI coordinates | | | | |
|---|---|---|---|---|---|---|---|
| **Region (Putative)** | **Laterality** | **Cluster Size** | **X** | **Y** | **Z** | **Max stat t** | **P Cluster Corrected** |
| Superior Frontal Gyrus/Middle Frontal Gyrus | Right | 245 | 26 | 58 | 14 | 6.9 | 0.002 |
| Superior Frontal Gyrus, Middle Frontal Gyrus (BA 6) | Right | 269 | 8 | -2 | 64 | 6.26 | 0.001 |
| Middle Temporal Gyrus | Right | 157 | 50 | -28 | -8 | 5.97 | 0.016 |
| Superior Frontal Gyrus/Middle Frontal Gyrus (BA 8, BA 9) | Right | 188 | 4 | 40 | 46 | 5.28 | 0.009 |
| Insula, Inferior Frontal Gyrus (BA 13, BA 47) | Left | 174 | -32 | 18 | -10 | 4.73 | 0.013 |
| Insula, Inferior Frontal Gyrus | Right | 313 | 44 | 22 | -6 | 4.69 | <0.001 |

Table S42: Negative loading on feedbackLoss – feedbackWin in money session (This is positive loading on feedbackWin – feedbackLoss in money session) (n = 20)

| | | | Peak MNI coordinates | | | | |
|---|---|---|---|---|---|---|---|
| Region (Putative) | Laterality | Cluster Size | X | Y | Z | Max stat t | P Cluster Corrected |
| Middle Temporal Gyrus/Angular Gyrus (BA 39) | Left | 151 | -44 | -72 | 30 | 6.11 | 0.025 |
| Posterior Cingulate, Lingual Gyrus (BA 18), Cerebellum_4_5_left (Culmen) | Left | 268 | -10 | -54 | 8 | 5.99 | 0.001 |
| Posterior Cingulate (BA 23) | | 180 | 8 | -52 | 8 | 5.13 | 0.011 |

#### GLM C:

##### Choice*ST regressor

No significant clusters were found with n = 20.
